# Cross-scale dynamics and the evolutionary emergence of infectious diseases

**DOI:** 10.1101/066688

**Authors:** Sebastian J. Schreiber, Ruian Ke, Claude Loverdo, Miran Park, Prianna Ahsan, James O. Lloyd-Smith

## Abstract

When emerging pathogens encounter new host species for which they are poorly adapted, they must evolve to escape extinction. Pathogens experience selection on traits at multiple scales, including replication rates within host individuals and transmissibility between hosts. We analyze a stochastic model linking pathogen growth and competition within individuals to transmission between individuals. Our analysis reveals a new factor, the cross-scale reproductive number of a mutant virion, that quantifies how quickly mutant strains increase in frequency when they initially appear in the infected host population. This cross-scale reproductive number combines with viral mutation rates, single-strain reproductive numbers, and transmission bottleneck width to determine the likelihood of evolutionary emergence, and whether evolution occurs swiftly or gradually within chains of transmission. We find that wider transmission bottlenecks facilitate emergence of pathogens with short-term infections, but hinder emergence of pathogens exhibiting cross-scale selective conflict and long-term infections. Our results provide a framework to advance the integration of laboratory, clinical and field data in the context of evolutionary theory, laying the foundation for a new generation of evidence-based risk assessment of emergence threats.

## Introduction

Emerging infectious diseases are rising in frequency and impact and are placing a growing burden on public health and world economies [1–4]. Nearly all of these emergence events involve pathogens that are exposed to novel environments such as zoonotic pathogens entering human populations from non-human animal reservoirs, or human pathogens exposed to antimicrobial drugs [1]. In these novel environments, pathogens may experience new selective forces acting at multiple biological scales, leading to reduced replication rates within hosts or less efficient transmission between hosts. When these novel environments are sufficiently harsh, emergence only occurs when the pathogen adapts sufficiently quickly to avoid extinction. As genetic sequencing of pathogens becomes increasingly widespread, there are clear signs of such rapid adaptation [5–11], but we lack a cohesive framework to understand how this process might work across scales. Theoretical studies have shed important insights into circumstances under which this evolutionary emergence is possible, but either have focused on the host-to-host transmission dynamics and treated within-host dynamics only implicitly [12–15], or have accounted for explicit within-host dynamics only along a fixed transmission chain [16, 17]. Here, we introduce and analyze a model explicitly linking these two biological scales and demonstrate how within-host viral competition, infection duration, transmission dynamics within a host population, and the size of transmission bottlenecks determine the likelihood of evolutionary emergence. This analysis sheds new light on factors governing pathogen emergence, addresses long-standing questions about evolutionary aspects of emergence, and lays the foundation for making risk assessments which integrate outcomes from *in vitro* and *in vivo* experiments with findings from sequence-based surveillance in the field.

Recent empirical findings have highlighted the need for a new generation of theory on pathogen emergence, which addresses the current frontiers of dynamics within hosts and across scales. For most pathogens, and certainly for RNA viruses and single-stranded DNA viruses, individual hosts often are not dominated by single pathogen genotypes [18, 19]. Furthermore, at the host population scale, pathogen allele frequencies at a given locus exhibit a range of dynamics from rapid selective sweeps for drug resistance or immune escape [20–22] to gradually changing frequencies [23, 24]. Together, these observations lead to the long-standing question of whether adaptive evolution of viruses occurs within single hosts by rapid fixation of beneficial mutants, or more slowly by a gradual shift of allele frequencies along chains of transmission [25, 26]. A recent wave of studies tracking changes in within-host genetic diversity through chains of transmission among hosts [27–34] provide unique opportunities to address this question, but a theoretical framework is needed.

Empirical studies, together with analyses at broader population scales, have highlighted the crucial influence of the transmission process – and particularly the population bottleneck associated with transmission – in filtering viral diversity. The existence of transmission bottlenecks has long been recognized, and is hypothesized to play a critical role in pathogen evolution [35–40]. Recent studies have reported that bottleneck widths vary considerably among pathogens and routes of transmission [41, 42], and perhaps across different phases of host adaptation [43]. Narrow transmission bottleneck sizes of 1 to 2 viral genotypes are common for HIV-1 [44–46] and hepatitis C virus [47, 48], and bottlenecks of 1 to 3 viruses are reported for influenza in ferret respiratory droplet transmission experiments [41, 42, 49] and in some studies of natural human transmission [50, 51]. Much wider bottleneck estimates, of 30 to over 100 viruses, have been reported for natural transmission of influenza in horses [29] and swine [52]; for ferret transmission experiments via direct contact [41, 42]; and for transmission of Ebola virus among humans [53]. While wide bottlenecks were also reported for natural influenza transmission among humans [54, 55], this was determined to be a bioinformatic artefact [56].

A major frontier in understanding viral adaptation is how the transmission process influences evolution at population scales. Past work has emphasized the potentially deleterious effect of genetic drift [35, 37, 39], but a rising tide of studies reports direct selection for transmissibility. This can arise as a strong selection bias at the transmission bottleneck, where strains present at low or undetectable frequencies in the donor host are preferentially transmitted to the recipient [43, 49, 57, 58], or it can be measured directly via experimental infection and transmission studies [24, 59–61] (though we emphasize that enhanced transmissibility is not inevitable, and depends on availability of suitable adaptive genotypes [62]). Overall transmission rates can be viewed as being determined by total viral loads, weighted by genotype-specific transmissibilities [58]. Importantly, the transmissibility trait can vary independently from viral replication fitness within hosts, so there is potential for conflicts in selection across scales. Indeed, there is clear evidence that HIV-1 has certain genotypes that transmit more efficiently, but then the within-host population tends to evolve toward lower-transmission strains during an infection [46, 58, 63–65]; a similar phenomenon has been reported for H5N1 influenza [49] and H9N2 influenza [66]. In an extreme example, <u>Plasmodium</u> parasites were found to rapidly evolve resistance to an antimalarial drug, but at the cost of complete loss of transmissibility [67]. Experimental evolution studies have highlighted how antagonistic pleiotropy can lead to tradeoffs between viral replication and the extracellular survival that is required for transmission [68, 69], and a similar tradeoff has been postulated for environmental transmission of avian influenza in the field [70]. Together these findings contribute to a growing evidence base that cross-scale conflicts in selection may inhibit the emergence of new viral strains in many systems [reviewed in 15]. Collectively these empirical findings highlight the need for a theory of evolutionary emergence that accounts explicitly for the within-host dynamics of competing viral strains, transmission bottlenecks, and host-to-host transmission dynamics [71]. To this end, we introduce and analyze a model which integrates previous work on stochastic models of evolutionary emergence and deterministic models explicitly coupling within- and between-host dynamics [12, 14, 16, 17, 60, 65, 72–75]. Our analysis allows us to address several fundamental questions about the emergence of novel pathogens: What factors limit evolutionary emergence for pathogens with different life histories? Why do some apparently ‘nearby’ adaptive mutants fail to emerge? How do bottleneck sizes influence the likelihood of emergence? Do evolutionary changes occur swiftly within individual hosts, or gradually across chains of transmission? Moreover, our analysis allows us to examine the relative importance of genetic diversity in zoonotic reservoirs versus the acquisition of new mutations following spillover into humans [76–78]. Specifically, we address the long-standing question of how much emergence risk is increased if the “spillover inoculum” includes some genotypes bearing adaptive mutations for the novel host? Finally, our analysis enables us to unify findings from previous theoretical studies, and propose mechanistic interpretations of phenomenological parameters from earlier work.

## Models and Methods

Our stochastic multiscale model of evolutionary emergence follows a finite number of individuals in a large, susceptible host population exposed to a pathogen from a reservoir population (Fig 1A). Although our framework represents many types of pathogens and can be extended to any number of strains, we focus on the case of a viral pathogen with two strains: a wild-type maladapted for the novel environment and a mutant strain potentially adapted for the novel environment.

**Figure 1.**
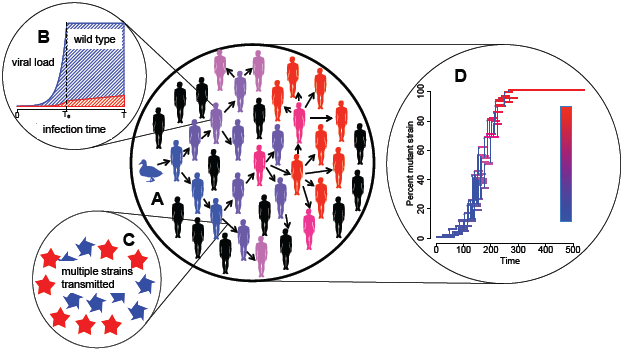
The cross-scale dynamic of evolutionary emergence. An individual is initially infected from the reservoir host population (panel A) with only the wild type viral strain (in blue). Within an infected individual (panel B), the viral load increases at an exponential rate until saturating at day *T*_*e*_ and ending after *T* days. Mutations ensure individuals typically have a mixed infection with wild-type (blue) and mutant (red) viral strains (panel B). The likelihood of transmission between individuals, and the composition of the infecting dose (panel C), depend on the size and composition of the infected individual’s viral load at the time of contact, and on the transmissibility of each strain. As the infection spreads in the population (panel A), the frequency of the mutant virions among infected individuals varies (panel D) and, ultimately, determines whether evolutionary emergence occurs. In D, each horizontal line marks the infectious period of an individual whose infection was initiated with that percentage of the mutant strain and the vertical arrows represent transmission events between individuals.

### The cross-scale model with explicit within-host dynamics

Formally, the cross-scale model is a continuous time, age-dependent, multi-type branching process [79, 80]. The “type” of individual corresponds to the composition of the initial virus population (i.e. the founding viral population that initiates the infection), and the “age” of an individual corresponds to the time since their initial infection. Within an infected host, the viral dynamics determine how the viral load and viral composition change over time due to competition between strains and mutation events. Transmission events are determined by the viral load and composition of the host and, consequently, are age-dependent. Below, we describe the model at each scale and the biological processes we consider in detail. The mean-field analogue of our model is an age-structured partial differential equation model introduced by Coombs et al. [73].

#### Within-host scale model

Infection of a host usually starts locally at the site of viral entry or first tissue contact. This local spread involves a small number of viruses, their infection of host cells at the exposure interface and possibly the innate immune response to infection [81, 82]. During this period, infection is stochastic and establishment of infection is not guaranteed [82, 83]. When one or more virions survive the period of initial local spread, they establish lineages that comprise the productive infection. These virions are termed as transmitted founder viruses [44]. Here, we explicitly define the number of transmitted founder viruses that establish productive infection as the bottleneck width *N*. This quantity can be estimated using viral genetic sequencing data, for example in [44, 55]. Our within-host model starts with the transmitted founder viruses by assuming an initial viral load *N*, and hence considers the period of established productive infection only (as with other within-host models [84, 85] and cross-scale deterministic models [73]). The initial viral exposure and stochastic local infection process are implicitly incorporated into the transmission term in the population scale model as described below, and consequently, we consider a successful transmission event as a transmission event leading to an established infection.

The within-host dynamics are modeled with coupled differential equations where *v*(*t*) = (*v*_*w*_(*t*), *v*_*m*_(*t*)) denotes the vector of viral abundances

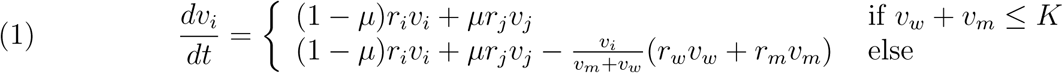

where *i* ≠ *j* are either *w* or *m*, for wild-type and mutant strains respectively, and *µ* is the mutation rate between these strains. At time *t* = 0, *v*(0) = (*v*_*w*_(0), *v*_*m*_(0)) corresponds to the initial viral load of an infected individual, and *v*_*w*_(0) + *v*_*m*_(0) = *N*. Our within-host model is similar in structure to the quasi-species model of Lythgoe et al. [65].

In this model, the viral population initially increases exponentially because of the availability of a large number of target cells. The wild-type and mutant strains increase exponentially at rates, *r*_*w*_ and *r*_*m*_, respectively. These dynamics are consistent with the viral dynamics predicted by standard multistrain within-host models when target cells are abundant [86–88]. The viral load saturates at time *T*_*e*_ with a maximal viral load *K* (Fig 1B). We assume that after *T*_*e*_, the viral population size stays constant at *K*, and the frequencies of the wild-type and the mutant change due to their fitness differences. We further assume that the infectious period starts when *v*_*w*_(*t*) + *v*_*m*_(*t*) *>* 0 and ends after *T* days. For some acute infections, viral load usually decreases rapidly after viral peak (e.g., influenza A infections [89]), and thus *T* is close to *T*_*e*_. For other acute infections, such as SARS-CoV-2 [90] infection, viral load remains at a high level after peak viremia for a longer period of time, in the range of weeks. In this case, *T > T*_*e*_. For chronic infections (though our work does not necessarily consider the full range of evolutionary processes involved in chronic infections) such as HIV and Hepatitis C [91] infections, viral loads usually reach a set-point and can stay roughly constant for an extended period of time, in the range of years. In this case, *T* is much greater than *T*_*e*_. Overall, this within-host model serves as a flexible framework to describe a range of viral dynamics from both acute and chronic infections, while maintaining simplicity to enable analysis.

#### Population scale model

At the scale of the host population, the transmission dynamics are modeled using a multi-type branching process. Each infectious individual encounters a Poisson-distributed number of susceptible individuals at a rate of *β* individuals per day. Then, the number of contacts of an infected individual during the infectious period is Poisson distributed with mean *βT*. Each contact results in a successful transmission event with probability *p*(*E*) where *E* is the effective viral load at the time *t* of transmission (see below). Similar to the deterministic model of Lythgoe et al. [65], *p*(*E*) is an increasing function of *E*. Our main analyses assume that the transmission function *p*(*E*) is linear, but nonlinear transmission functions yield nearly identical results (Figs. Appendix–2 through 4).

The effective viral load *E* is calculated as *E* = *b*_*w*_*v*_*w*_(*t*) + *b*_*m*_*v*_*m*_(*t*), where *b*_*w*_ and *b*_*m*_ are the transmissibilities of the wild-type and the mutant, respectively. Here *b*_*w*_ and *b*_*m*_ account for the survival of the viruses through a range of processes during transmission, including their likelihoods of being shed from the donor host, surviving the environment outside of a host, and reaching and infecting the target cells in an uninfected host. Furthermore, as explained in the within-host model section, these parameters also account for the likelihoods of the viruses to survive initial local infection and establish a productive infection in the recipient host. Viruses may face different challenges and selection pressures to overcome the barriers in each of these processes. Here, for simplicity and generality, a single parameter is used to summarize the transmissibility of different viruses because little is known or measured about the ability of a virus to overcome these barriers. More explicit models can be constructed as the relevant data become available.

In the event of successful transmission, there are *N* virions (the transmitted founder viruses) that establish the productive infection. In the model, these *N* virions are sampled binomially from the source individual’s viral load weighted by the transmissibilities of the viral strains. Thus, the probability of infecting an individual with a viral load of 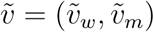 with 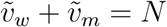 equals

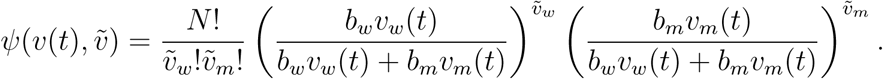

Under these assumptions, during their infectious period, an infected individual of type *v*(0) infects a Poisson distributed number of individuals with viral load 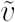, and the mean of this distribution equals

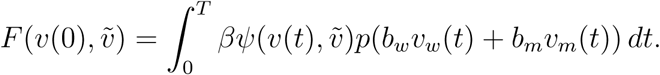

Overall, by explicitly modeling the cross-scale dynamics, our model simultaneously tracks the number of infected hosts and the viral loads within each infected host (Fig 1D). The structure of our stochastic model is similar to Peck et al. [16]’s stochastic model of molecular viral evolution along transmission chains. However, our model accounts for stochastic transmission dynamics rather than conditioning on a chain of transmission, and explicitly accounts for the dynamics of competing viral strains. It also has similarities with Geoghegan et al. [17]’s cross-scale, stochastic model of a single transmission event from a donor host to a recipient host. Like our model, Geoghegan et al. [17]’s model has constant transmission bottlenecks, multinomial sampling from donor to host, and explicit within host dynamics with exponential growth and ceiling phases. Their model, however, focuses on a single transmission event and assumes that all virions are equally likely to be transmitted from donor to host, i.e. it assumes no selection based on transmissibility.

## Methods

To solve the probabilities of emergence, we use the discrete-time branching process given by censusing the infected population at the beginning of each generation of infection. This discrete-time process is known as the embedded process [79, 80]. All the statistics of this embedded process are given by the probability generating map *G* : [0, 1]^*N*+1^ → [0, 1]^*N*+1^ where *N* + 1 is the number of types of initial viral loads i.e. all combinations of wild-type and mutant virions for *N* virions [79, 80]. We index the coordinates by the initial number of mutant virions 0, 1, 2, …, *N* within an infected individual and have

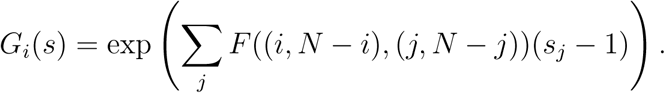

The *i*-th coordinate of

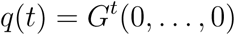

is the probability of extinction by generation *t* when there is initially one infected individual with initial viral load (*i, N* − *i*). The probability of emergence is given by 1 −*q* where *q* = lim_*t*→∞_ *q*(*t*) is the asymptotic extinction probability. The limit theorem of branching processes implies that *q* = (*q*_0_, …, *q*_*N*_) is the smallest (with respect to the standard ordering of the positive cone), non-negative solution of the equation *q* = *G*(0, …, 0). These extinction probabilities can be non-zero if and only if the dominant eigenvalue of the Jacobian matrix *DG*(1, 1, …, 1) is greater than one. Equivalently, the reproductive number given by the dominant eigenvalue of the next generation matrix of *DG*(1, 1, …, 1) is greater than one [92]. Note that the linear map *s ↦ DG*(1, 1, …, 1)*s* corresponds to the mean-field dynamics of the embedded multi-type branching process.

For the numerical work, we used linear, logarithmic, and saturating functions for the transmission probability function *p*. All gave similar results but we present the linear case as most analytical results were derived for this case. To compute the asymptotic extinction probabilities, we iterated the probability generating map *G* for 2, 000 generations. For the individual based simulations, we solved the within-host differential equations using matrix exponentials and renormalizing these exponentials when the viral load reached the value *K*. Between host transmission events were determined by a time-dependent Poisson process with rate function *p*(*b*_*w*_*v*_*w*_(*t*) + *b*_*m*_*v*_*m*_(*t*)), and multinomial sampling was used to determine the initial viral load of an infected individual.

## Results

### The probability of evolutionary emergence

We first focus on the scenario of a single individual in the host population getting infected by the wild-type strain. We assume that the mean number of individuals infected by this individual (the reproductive number *R*_*w*_ of the wild-type) is less than one. Hence, in the absence of mutations, there is no chance of a major outbreak [92]. However, if the wild-type strain produces a mutant strain whose reproductive number *R*_*m*_ is greater than one, there is a chance for a major outbreak. The mutant strain might have a higher reproductive number than the wild-type strain because it replicates more rapidly within the host or because it transmits more effectively to new hosts (or both). We define these within-host and between-host selective advantages as *s* = *r*_*m*_ − *r*_*w*_ and *τ* = log(*b*_*m*_) − log(*b*_*w*_), respectively.

Consistent with theoretical expectations, a non-zero probability of evolutionary emergence requires the mutant’s reproductive number *R*_*m*_ to be greater than one (Fig 2). However, the mixture of selective advantages or disadvantages of the mutant strain that give rise to *R*_*m*_ *>* 1 depends in a complex manner on the pathogen’s life history traits, such as the duration of the infection (Fig 2A,B vs. C,D) and the transmission bottleneck width (Fig 2A,C vs. B,D). Notably, for long-term infections with a large transmission bottleneck size, the emergence probability can be effectively zero (i.e. *<* 10^−16^) for mutant strains whose reproductive number exceeds one (white region bounded by the solid red curve in Fig 2D) – a finding not explained by classical theory. As we shall show, this outcome is predicted by a new quantity, the cross-scale reproductive number (*α*) of a mutant virion – the mean number of mutant virions transmitted to susceptible individuals by an infected individual whose initial viral load contains 1 mutant virion and *N* − 1 wild type virions. When *α* is less than one, evolutionary emergence becomes unlikely.

**Figure 2.**
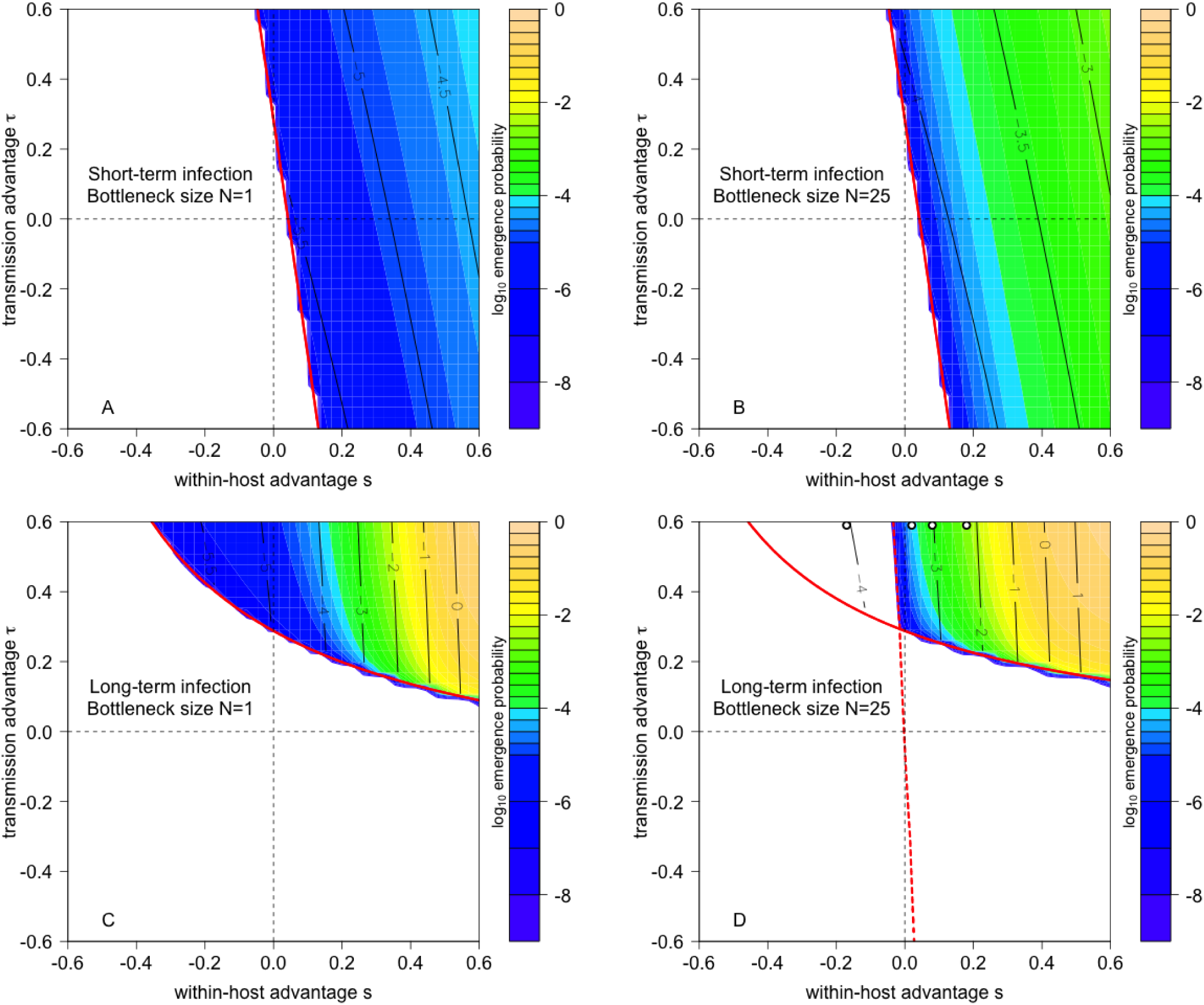
The joint effects of within-host and between-host selective advantage of the mutant on the probability of emergence for pathogens with short-term (A,B) and long-term (C,D) infectious periods, and with transmission bottlenecks of size *N* = 1 (A,C) and *N* = 25 (B,D). Colorations correspond to log_10_ of the emergence probability. The critical value of the mutant reproductive number *R*_*m*_ equaling one is drawn in solid red. Black contour lines correspond to log_10_ of the number of mutant virions transmitted by an individual initially only infected with the wild strain. In D, the critical value of the cross-scale reproductive number, *α* = 1, of a mutant virion is shown as a red dashed line and the white circles correspond to the *τ* and *s* values used in Fig 3. Parameters: *K* = 10^7^, *βT* = 30, *T* = 7.5 *<* 12.9 = *T*_*e*_ (short-term infection) and *T* = 30 *>* 12.9 = *T*_*e*_ (long-term infection), *b*_*w*_ chosen so that *R*_0_ = 0.75 for wild type, *r*_*w*_ = 1.25 and *µ* = 10^−7^.

To understand these complexities, we determine the conditions under which the mutant’s reproductive number *R*_*m*_ exceeds one, and then present analytic approximations for the emergence probability when *R*_*m*_ *>* 1.

### Cross-scale selection and the mutant reproductive number

*R*_*m*_. The reproductive numbers of the wild-type strain (*R*_*w*_) and mutant strain (*R*_*m*_) can be calculated as the product of the contact rate, the average transmissibility of the strain during the infectious period, and the infection duration (see, Coombs et al. [73] and Appendix). These reproductive numbers are positively correlated with the contact rate, infection duration, transmissibility, and viral per-capita growth rates. The influence of the maximal viral load *K* depends on the infection duration. For short-term infections, defined here as infections with a relatively short saturated phase (i.e. *T* − *T*_*e*_ ≪ *T*_*e*_), increasing *K* has little effect on a strain’s reproductive number. For long-term infections, defined here as as infections with a long saturated phase (i.e. *T* − *T*_*e*_ ≫ *T*_*e*_), reproductive numbers increase with *K*.

Whether a selective advantage at either scale results in the mutant reproductive number *R*_*m*_ exceeding one depends on the duration of the infection. For short-term infections, most transmissions occur towards the end of the infectious period *T*, when the infectious load is the highest. By the end of the infectious period, the mutant viral density has increased approximately by a factor of *e*^*sT*^ more than the wild-type, and transmission for each mutant virion is exp(*τ*) more likely than for a wild-type virion. Refining this intuition, we derive the approximation (Appendix)

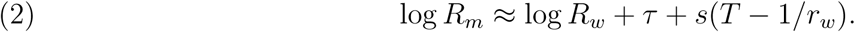

This approximation shows that a sufficiently strong selective advantage at either scale can result in the mutant reproductive number exceeding one (*R*_*m*_ *>* 1) despite a selective disadvantage at the other scale (confirmed by exact calculations in Fig 2A,B). For short-term infections where viral dynamics are dominated by the exponential phase, the longer the duration of infection, the greater the influence of the within-host selective advantage compared to the between-host selective advantage (e.g., steep contours in Fig 2A).

For long-term infections, the viral load will tend to *K* for both purely wild-type or purely mutant infections. Thus, the only difference will be in transmissibility and we get the approximation (Appendix)

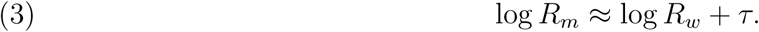

This approximation implies that a between-host selective advantage is required for a long-term infection to be capable of evolutionary emergence (confirmed by exact calculations in Fig 2C). When viral dynamics are dominated by the saturated phase at fixed *K*, a within-host selective advantage has little impact on the average viral load during the infectious period of an individual solely infected with the mutant strain and, consequently, provides a minimal increase in the mutant reproductive number.

### Going beyond the mutant reproductive number

When the mutant strain has a reproductive number greater than one, there is a non-zero probability of a major outbreak that is well-approximated by the product of three terms (compare Fig. 2 to Fig. Appendix–6):

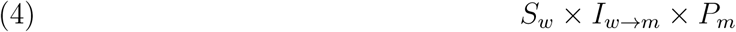

where *S*_*w*_ is the mean size of a minor outbreak due to the wild type, *I*_*w*→*m*_ is the mean number of individuals infected with a mutant virion by an individual initially only infected with the wild type, and *P*_*m*_ is the probability an individual infected with one mutant virion causes a major outbreak. The magnitude of the probability *P*_*m*_ depends on the mutant reproductive number, *R*_*m*_, as in previous theory; however we show below that it is also determined strongly by a new quantity, the cross-scale reproductive number *α* of a mutant virion. Our approximation (4), which can be viewed as a multiscale extension of earlier theory [12, 13], highlights three key ingredients, in addition to *R*_*m*_ *>* 1, for evolutionary emergence.

First, the size of the minor outbreak produced by the wild type determines the number of opportunities for the mutant strain to appear within a host. The average size of this minor outbreak equals 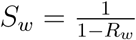, as noted by earlier studies [12, 13]. If the wild strain is badly maladapted, then it is expected not to spread to multiple individuals (e.g. if *R*_*w*_ *<* 1*/*2, then 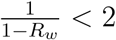) and opportunities for transmission of mutant virions are very limited. Alternatively, if the wild strain is only slightly maladapted to the new host, then, even without any mutations, the pathogen is expected to spread to many individuals (e.g. if *R*_*w*_ = 0.95, then 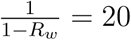), thereby providing greater opportunities for evolutionary emergence. Our analysis implies that higher contact rates, within-host viral growth rates, viral transmissibility, and maximal viral loads (for long infectious periods) facilitate these larger reproductive values.

Second, the mutant strain must be transmitted successfully to susceptible individuals — the second term *I*_*w*→*m*_ of our approximation (4). For an individual initially infected only with the wild-type strain, *I*_*w*→*m*_ equals the product of the contact rate, the infection duration, and the likelihood that a mutant virion is transmitted during a contact event, averaged over the full course of infection (Appendix). The likelihood of transmitting mutant virions on the *t*^*th*^ day of infection is proportional to the product of the transmission bottleneck width (*N*), the within-host frequency of the mutant strain, and the transmissibility *b*_*m*_ of the mutant strain. This highlights an important distinction between short-term and long-term infections. For short-term infections where *sT* is small, there is insufficient time for the frequency of mutants to rise within a host, so transmission events with mutant virions are rare (*<* 1*/*1, 000 for all black contour lines in Fig 2A,B). This is a key obstacle to evolutionary emergence in short-term infections. In contrast, for long-term infections where the mutant strain has a substantial within-host selective advantage, the mutant strain is transmitted frequently (e.g. the expected number of events *>* 1 for some contours in Fig 2C,D).

Finally, even if the mutant strain is successfully transmitted, an individual infected with the mutant strain needs to give rise to a major outbreak which occurs with probability *P*_*m*_, see equation (4). This requires the mutant strain to rise in frequency in the infected host population. A mean field analysis for larger bottleneck sizes (*N >* 5 in the simulations) reveals that mutant frequency initially grows geometrically by the cross-scale reproductive number *α* of a mutant virion–the number of mutant virions, on average, transmitted by an individual initially infected with a single mutant virion and *N* −1 wild type virions (Appendix). If this cross-scale reproductive number is greater than one, then each mutant virion replaces itself with more than one mutant virion in the next generation of infection, and the frequency of mutant virions increases in the infected host population. If the cross-scale reproductive number *α* is less than one, the frequency of mutants decreases, thereby hindering evolutionary emergence.

For short-term infections, the cross-scale reproductive number *α* is equal to the ratio of the basic reproductive numbers:

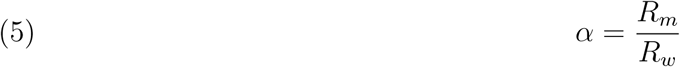

Thus for short-term infections there is no additional condition required for emergence. Whenever the mutant reproductive number *R*_*m*_ exceeds one, there is a mean tendency for the mutant strain to increase in frequency once it has been successfully transmitted to susceptible individuals (i.e. *α >* 1 because *R*_*m*_ *>* 1 *> R*_*w*_). The greater the ratio *R*_*m*_*/R*_*w*_, the more rapid the increase in frequency.

For long-term infections, there is sufficient time for within-host selection to change the frequency of the mutant strain within a host. Larger transmission bottlenecks increase the likelihood that these changes in frequency are transmitted between hosts. For these long infectious periods and larger bottlenecks, a within-host selective disadvantage reduces the cross-scale reproductive number *α* (Appendix):

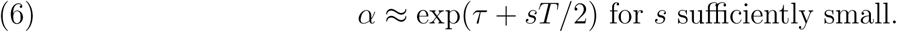

Hence, the cross-scale reproductive number *α* may be less than one even when the mutant reproductive number *R*_*m*_ is greater than one. This phenomenon, which arises from the interplay of dynamics at within-host and between-host scales, moderated by the transmission bottleneck width, explains the puzzling behavior about the emergence probabilities noted earlier (the white region bounded by solid and dashed red lines in Fig 2D).

The importance of these frequency dynamics can be visualized via individual-based outbreak simulations, and cobwebbing diagrams summarizing the mean field dynamics. When the mutant reproductive number *R*_*m*_ is greater than one but its cross-scale reproductive number *α* is less than one, mutant virions may be transmitted but the resulting mixed infections are invariably taken over by purely wild-type infections (Fig 3A). Only pure mutant infections can escape this “relapse” to wild-type, and then only if the mutation rate *µ* is low enough that new wild-type virions are very slow to appear. When the within-host selective disadvantage is weak and the between-host selective advantage is strong, the cross-scale reproductive number *α* may be slightly greater than one and the mutant strain can drift to higher frequencies within the infected host population (Fig 3B). For large within-host selective advantages, the cross-scale reproductive number *α* is large and the mutant strain can sweep rapidly to fixation in the infected host population (Fig 3C). Thus, in addition to revealing a new condition needed for evolutionary emergence, the cross-scale reproductive number *α* summarizes the conditions under which evolution occurs swiftly or gradually within chains of transmission.

**Figure 3.**
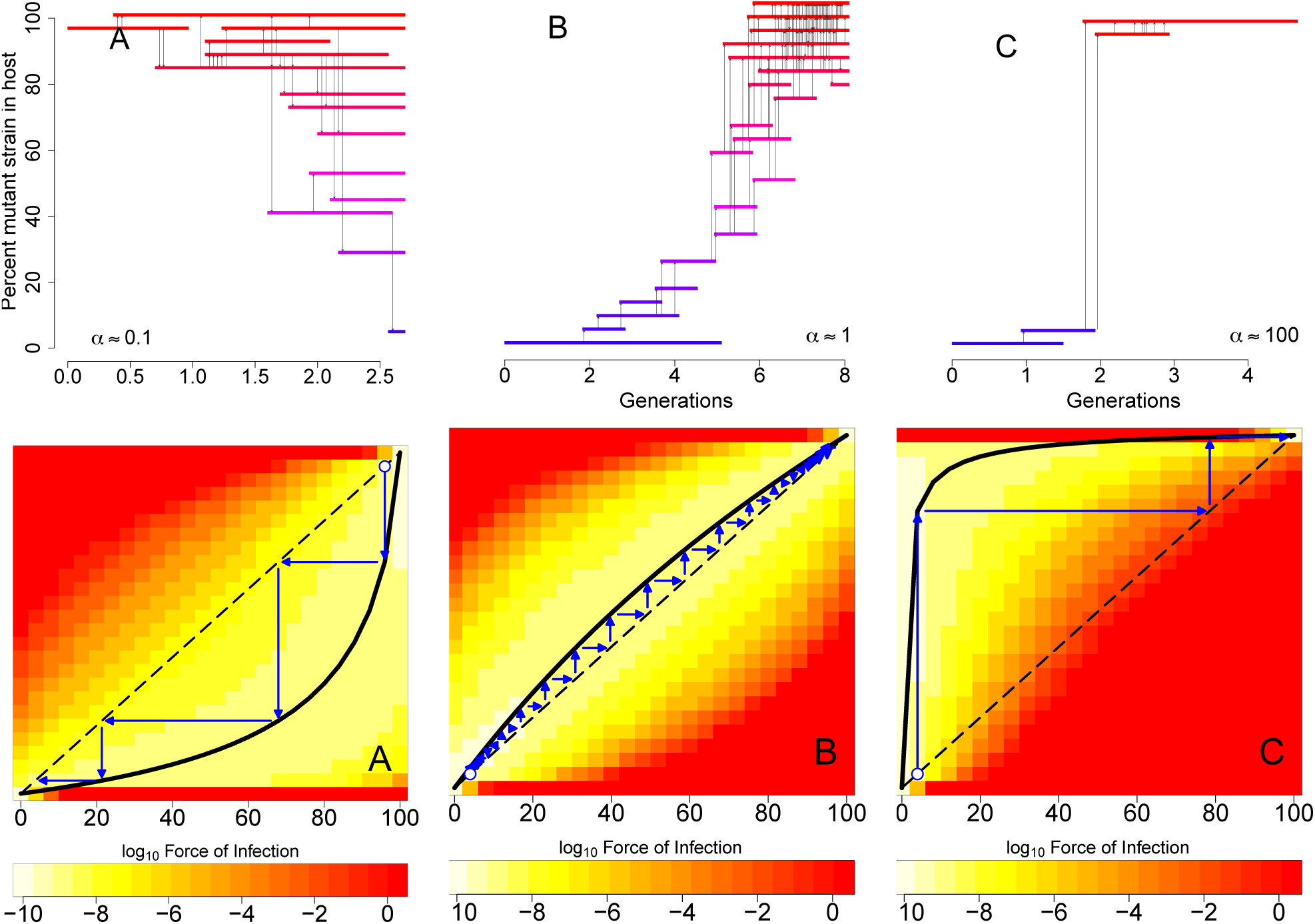
Frequency dynamics of the mutant strain in the host population. For long-term infections with moderate to large transmission bottlenecks (*N >* 5), individual-based simulations corresponding to three values of the cross-scale reproductive number *α* of a mutant virion illustrate (A) the mutant strain decreasing in frequency (despite an index case initially only infected with the mutant strain) when the cross-scale reproductive number *α* is less than one, (B) a gradual sweep to fixation of the mutant strain when *α* ≈ 1, and (C) fast sweeps to fixation for large values of *α >* 1. In these individual based simulations, each horizontal line marks the infectious period of an individual whose infection was initiated with that percentage of the mutant strain and the vertical arrows represent transmission events between individuals. In the bottom half of the figure, the mean field dynamics corresponding to each of the individual-based simulations are plotted as cobwebbing diagrams. The solid black curves correspond to the expected frequency of the mutant strain in the infected host population in the next generation given the frequency in the current generation. Thin blue lines indicate how the expected frequencies change across multiple generations. The colored backgrounds represent the expected number of individuals infected with a certain percentage of the mutant strain (vertical axis) by an individual with an initial percentage of the mutant strain (horizontal axis). Parameter values as in Fig 2D indicated with black asterisks.

### The dueling effects of transmission bottlenecks

Wider bottlenecks increase the likelihood of evolutionary emergence for pathogens with a short infectious period, but can hinder or facilitate evolutionary emergence of long-term infections (Fig 4A,B). For short-term infections, evolutionary emergence is constrained primarily by the transmission of mutant virions by individuals initially infected with only the wild strain. Wider transmission bottlenecks alleviate this constraint, especially when the mutant strain is expected to increase rapidly within the infected population (*α* ≫ 1; Fig 4A). When the mutant strain rises slowly in the infected host population (*α* slightly greater than one), the emergence probability is insensitive to the bottleneck size, regardless of infection duration.

**Figure 4.**
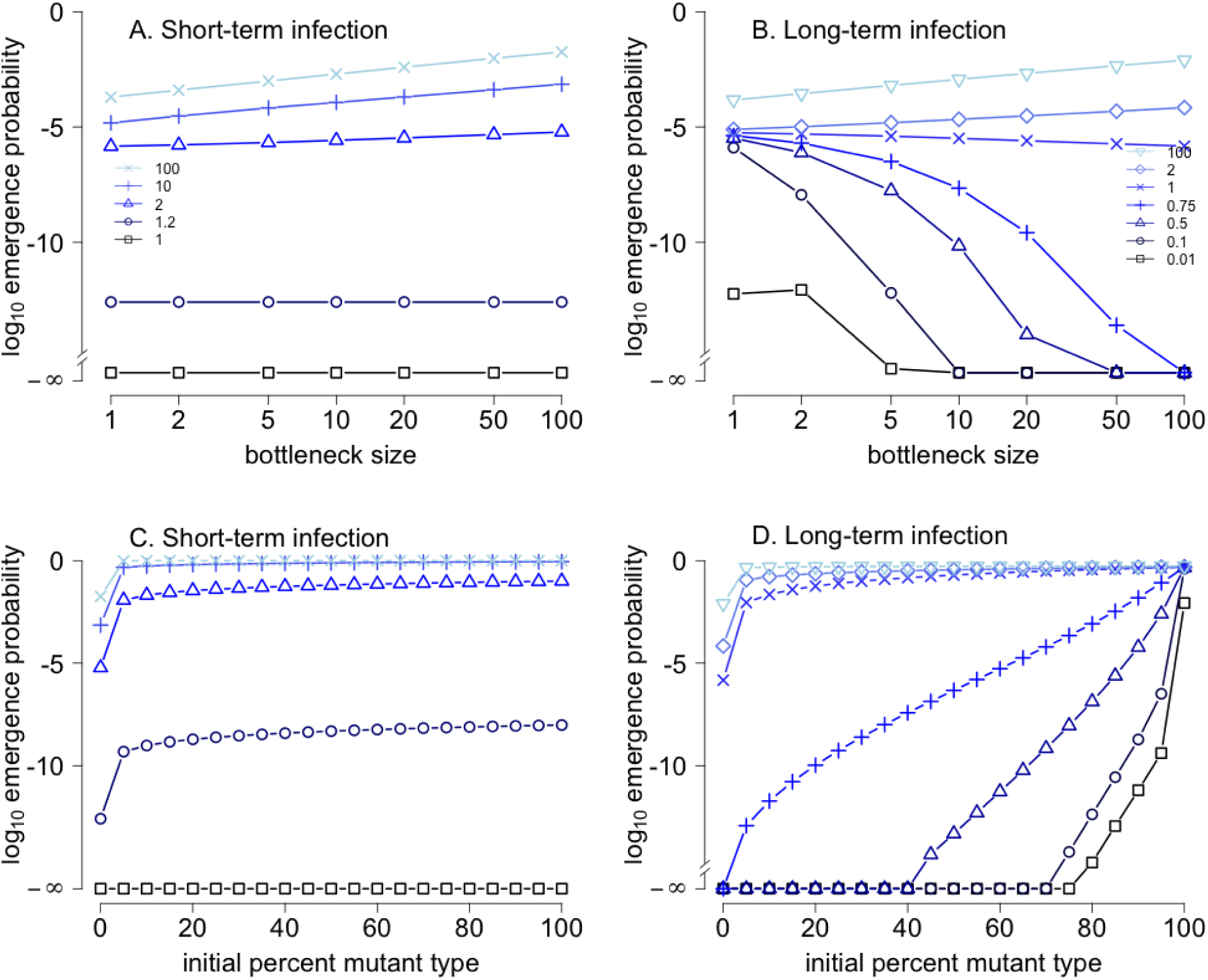
Effects of bottleneck size and mixed infections of the index case on evolutionary emergence of short-term and long-term infections. Different curves correspond to different tendencies, as measured by the cross-scale reproductive number *α*, for the mutant strain to increase in frequency in the infected host population. In (A) and (B), bottleneck size has negative effect on emergence when the cross-scale reproductive number *α* is less than one and a positive effect when *α* is greater than one. In (C) and (D), index cases initially infected with higher percentages of the mutant strain are more likely to lead to emergence. −∞ corresponds to numerical values of 10^−16^ or smaller. Parameters: *K* = 10^7^, *βT* = 150, *T* = 7.5 for short-term infections and *T* = 30 for long-term infections, *b*_*w*_ chosen so that *R*_0_ = 0.75 for the wild strain, *r*_*w*_ = 1.25, *τ* = 1, *s* chosen to achieve the *α* values reported in the legend, and *µ* = 10^−7^. *N* = 25 in (C) and (D).

For long-term infections for which the mutant strain’s reproductive number *R*_*m*_ is greater than one, but the cross-scale reproductive number *α* is less than one, emergence probabilities decrease sharply with bottleneck size (Fig 4B and Appendix). Because a mutant reproductive number *R*_*m*_ greater than one requires a between-host selective advantage (*τ >* 0) for a long-term infection, the cross-scale reproductive number *α* is less than one only if there is a within-host selective disadvantage (*s <* 0) so that mixed infections tend to be taken over by the wild-type. Consequently, the mutant virus can start an epidemic only when a host is infected with an initial viral load composed of mutant particles only, an event that becomes increasingly unlikely for larger bottleneck sizes *N*.

### Mutant spillover events hasten evolutionary emergence

When the mutant strain is circulating in the reservoir, the index case can begin with a mixed infection which invariably makes evolutionary emergence more likely (Fig 4C,D). For short-term infections, spillover doses that contain low or high frequencies of mutants have a roughly equal impact on emergence, and the magnitudes of these increases are relatively independent of the cross-scale reproductive number *α* (Fig 4C). This arises because the initial production and transmission of the mutant strain is the primary constraint on evolutionary emergence for short-term infections with *R*_*m*_ *>* 1 (black contours in Fig 2A,B). Consequently, mutant spillover events of any size are sufficient to overcome this constraint.

For long-term infections, the impact of mutant spillover depends on the cross-scale reproductive number *α*. When *α* is less than one, only spillover doses with high frequencies of mutants have a significant effect on emergence (i.e. bottom three curves in Fig 4D). When the cross-scale reproductive number *α* is greater than one, the effect mimics short-term infections and mutant spillover events of any size can substantially increase the chance of emergence (top three curves in Fig 4C,D).

## Discussion

We presented a cross-scale model for evolutionary emergence of novel pathogens, linking explicit representations of viral growth and competition within host individuals to viral transmission between individuals. Our work identifies four steps to evolutionary emergence summarized in Figure 5 and four ingredients (see, equation (4)) that govern these steps: (i) the reproductive number of the wild type which determines the size of a minor outbreak of this strain, (ii) the rate at which individuals infected initially with the wild-type strain transmit the mutant strain, and (iii) the cross-scale reproductive number *α* of a mutant virion which corresponds to the mean number of mutant virions transmitted by an individual whose initial infection only included one mutant virion, and (iv) the reproductive number of the mutant strain. Prior studies of evolutionary emergence [12–15] identified the importance of the single strain reproductive numbers, *R*_*m*_ and *R*_*w*_, and a phenomenological ‘mutation rate’, but ingredients (iii)–(iv) are new mechanistic insights arising from the cross-scale dynamics. By analyzing these ingredients of evolutionary emergence, we show how the probability of emergence is governed by selection pressures at within-host and between-host scales, the width of the transmission bottleneck, and the infection duration. We also map the conditions under which different broad-scale patterns are observed, from rapid selective sweeps to slower diffusion of new types. While our study has focused on within-host and between-host scales of selection, it could be generalized readily to other types of cross-scale dynamics where selection may act differently at different scales, such as within-farm and between-farm scales where genetic data have given insights into the emergence of high-pathogenicity avian influenza strains [93].

**Figure 5.**
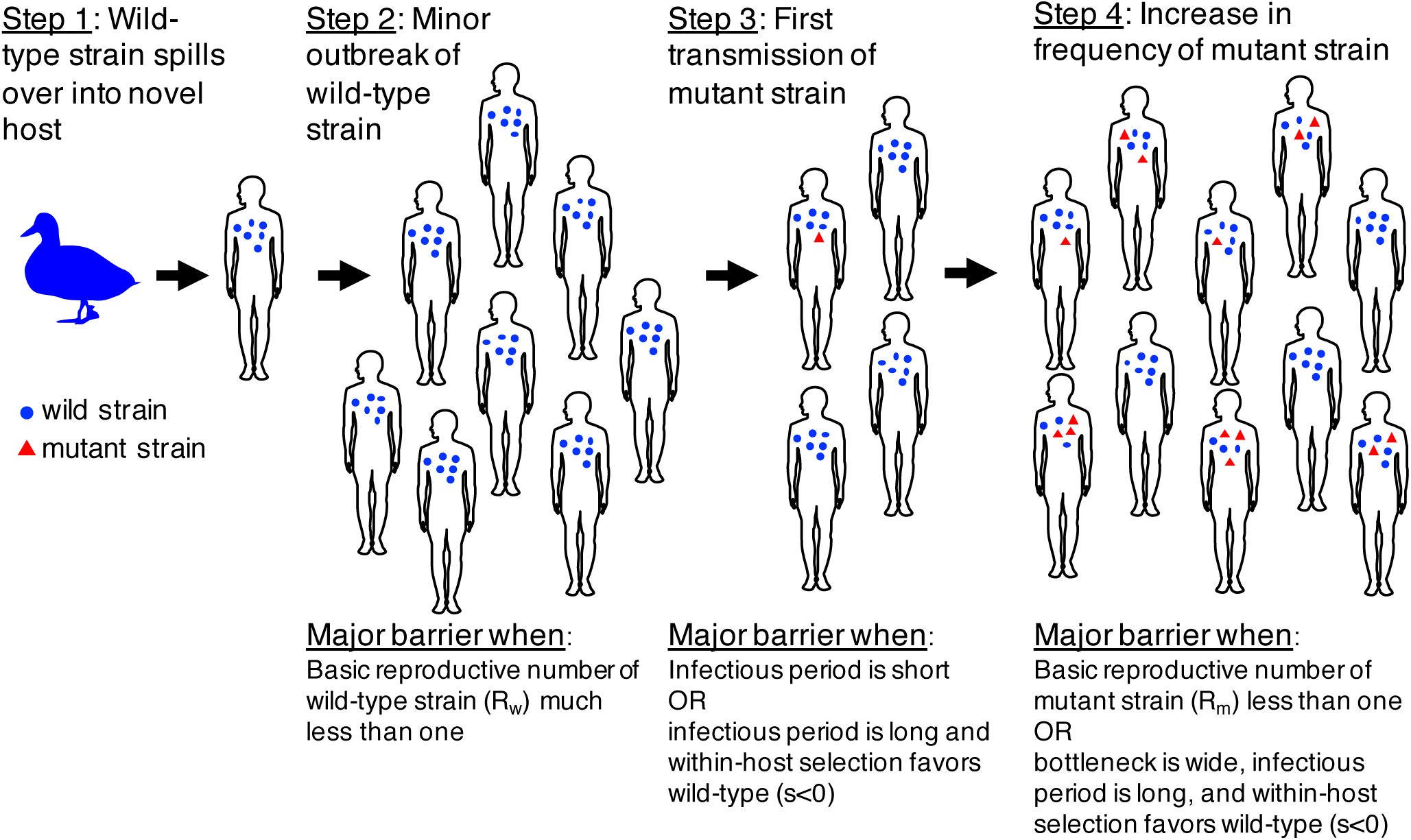
The major steps and barriers for evolutionary emergence.

Previous theoretical studies of evolutionary emergence of novel pathogens [12–15] have assumed infected individuals are, at any point in time, infected primarily by a single pathogen strain. Consequently, shifts from infection with one strain to infection with another must occur abruptly, relative to other processes. Such abrupt shifts could correspond to within-host selective sweeps or, if mutant strains remain at low frequency, to rare events in which only the mutant strain is transmitted. The seminal studies [12, 13] showed that under these conditions the probability of emergence is proportional to the frequency of these events, which they bundled together into a phenomenological “mutation rate”.

Our cross-scale analysis identifies the mechanistic counterpart to this phenomenological “mutation rate”, which is the probability that an individual infected initially with the wild-type strain ends up transmitting at least one virion of the mutant strain (Step 3 in Fig 5). This quantity, which is approximated by the black contours in Fig 2, is governed chiefly by the ability of the mutant strain to reach an appreciable frequency within the host over the course of an infection. This is evident from the strong dependence on the strength of within-host selection—which surprisingly is much stronger than the dependence on the transmission advantage of mutant virions—and the higher values found for larger bottleneck widths, which favor transmission of low-frequency mutants through a straight-forward sampling effect. This sampling effect is consistent with the theoretical work of Geoghegan et al. [17], and the experimental study of Frise et al. [42], who found larger bottlenecks increased the likelihood of mutant viral strains being transmitted between hosts. The duration of infection plays a crucial role, and our analysis showed that achieving this first transmission of the adaptive mutant is a key barrier to evolutionary emergence for short-term infections (Fig 2A,B). This finding aligns with the recent observation that putative immune-escape mutants of pandemic H1N1 influenza, which should have a within-host fitness advantage, were generated readily in infected humans but did not reach high within-host frequency and have been detected very rarely at the consensus level (i.e. they have failed to emerge) [94]. While more investigation is needed to determine the relevant *s* and *τ* parameters for these strains, these data are consistent with the mechanism we identify whereby these variants may be adaptive but have insufficient time to reach high enough frequencies to avoid being lost in transmission bottlenecks.

Our analysis highlights an additional factor, the cross-scale reproductive number *α* of a mutant virion, previously unrecognized in models neglecting within-host diversity and analyses centered on *R*_0_ for pure infections. Even after the mutant strain has been transmitted, it needs to increase in frequency at the scale of the infected host population (Step 4 in Fig 5). Specifically, each transmitted mutant virion, on average, needs to replace itself with more than one transmitted mutant virion in the next generation of infected hosts. When this occurs, it sets up a positive feedback along chains of infections: individuals with a higher frequency of the mutant strain tend to infect more individuals, which in turn provides more opportunities to transmit, on average, higher frequencies of the mutant strain to the next generation. Conversely, when this between-generation cross-scale reproductive number *α* is less than one, the positive feedback leads to lower and lower frequencies of the mutant strain within the infected host population. This positive feedback mechanism is stronger for wider transmission bottlenecks (*≥*5 virions in our numerical explorations), which better preserve the mutant frequency from one host to the next. Interestingly, this 5 virion threshold to define a wider transmission bottleneck is consistent with an earlier modeling study, which found that bottleneck sizes above 5 virions eliminated fitness losses in serial transfers of RNA viruses between cell culture plates [40].

The directionality of the positive feedback is more complex, and depends on multiple factors including the infection duration and the presence or absence of cross-scale conflicts. For long-term infections, mutant frequencies can drop deterministically within a host, and hence prevent emergence, even if the mutant strain has a reproductive number greater than one. This occurs when the mutant strain has a within-host selective disadvantage and between-host selective advantage (upper left quadrant of Fig 2D); the long infectious period allows time for the within-host disadvantage to drive the mutant strain to lower frequency and, thereby, set up the positive feedback effectively preventing evolutionary emergence. In contrast, for short-term infections the mutant strain tends to rise in frequency whenever the mutant reproductive number is greater than one, because there is insufficient time for any within-host disadvantage to act. In particular, evolutionary emergence may occur despite within-host selective disadvantages, a possibility excluded by previous theory [15]. Collectively these two results imply that, in the face of cross-scale conflict and wide transmission bottlenecks, longer infectious periods can inhibit, rather than facilitate [14], evolutionary emergence (Fig 2B,D). Related to this result, Geoghegan et al. [17] found that longer durations of the infectious period would lower the probability that a donor would infect the recipient. In their case, this occurred because fitness of the mutant was assumed to be lower in the donor host species and higher in the recipient species. Hence, a longer infectious period could purge any mutants arising in the donor and result in the recipient only receiving wild-type virions.

Previous theoretical studies examining the evolutionary consequences of cross-scale conflict [e.g. 65, 72, 73] differ from ours in several ways. Notably, they consider longer-term evolution for endemic diseases using deterministic models, rather than the inherently stochastic, shorter-term dynamics of evolutionary emergence. Using multiscale endemic SIR models, Coombs et al. [73] found that pathogen strains competitively superior at the within-host scale could be displaced by competitively inferior strains that had higher reproductive numbers at the epidemiological scale. This phenomenon was driven by non-equilibrium within-host dynamics, where early fluctuations in strain frequencies could have disproportionate influence if host survival was short. Our work reveals the converse case, where strains with lower reproductive numbers at the epidemiological scale (in fact, less than one) can prevent evolutionary emergence if they have a within-host advantage, by causing the adapted strains to have a cross-scale reproductive number *α* of less than one. Consistent with our result, Lythgoe et al. [65] showed found that deterministic, multistrain models could produce equilibrium states dominated by strains that were competitively superior at the within-host scale, despite reducing the reproductive number at the epidemiological scale. Parallel to our finding that cross-scale conflict occurred only for long-term infections, Lythgoe et al. [65]’s short-sighted evolution was most pronounced when within-host dynamics occurred at a faster time-scale.

Our cross-scale analysis also enables us to address two long-standing and interrelated questions in emerging pathogen research, regarding the influence of transmission bottleneck size on emergence probability and the importance of “pre-adapted” mutations circulating in the animal reservoir [17, 71, 76–78, 95]. In both cases, the answer depends on the cross-scale reproductive number *α* of a mutant virion that governs the frequency feedback. Under most circumstances, wider bottlenecks boost the probability of emergence (Fig 4A,B), because they favor the onward transmission of mutant virions when they are rare; this is particularly vital for the first transmission of mutant virions (i.e. Step 3 in Fig. 5). The exception is for long-term infections with *α <* 1, such that the mutant tends to decline in frequency in the infected host population. Under these circumstances, wider bottlenecks hinder emergence by propagating reductions in the frequency of the mutant strain more efficiently from host to host (Step 4 in Fig. 5). Conventional thinking about the influence of bottlenecks on viral adaptation emphasizes fitness losses due to genetic drift and the effects of Muller’s ratchet [35–37, 39, 40], which become more severe for narrower bottlenecks. Contrary to these negative effects of narrow bottlenecks, our findings highlight that narrower bottlenecks can aid emergence in long-term infections with a cross-scale conflict in selection (Fig 4B). Here the adaptive gain in transmissibility at population scales can be impeded by the selective disadvantage at the within-host scale, but, intriguingly, this disadvantage is neutralized by genetic drift arising from narrow bottlenecks. Given the evidence for cross-scale evolutionary conflicts for HIV-1 [58, 63, 64, 96], our results suggest the possibility that HIV-1’s narrow transmission bottleneck [44] could play a role in the emergence of novel strains (e.g. drug resistant strains).

Similar mechanisms dictate the influence of mutant viral strains circulating in the reservoir, particularly for long-term infections (Fig 4C,D). If the cross-scale reproductive number *α* of a mutant virion is greater than one, so that the mutant frequency rises easily in the infected host population, then even low frequencies of mutants in the reservoir lead to substantial risk of emergence. Indeed, for long-term infections with *α >* 1, emergence becomes almost certain when there are mutants in the initial spillover inoculum. Conversely, when the cross-scale reproductive number *α* is less than one, emergence probability scales with the proportion of mutants in the initial dose, and when *α* ≪ 1, the initial dose must consist almost entirely of the mutant strain in order to pose any major risk. These findings yield direct lessons for the growing enterprise of conducting genetic surveillance on zoonotic pathogens in their animal reservoirs [97–100]. A crucial requirement for effective genetic surveillance is the ability to identify genotypes of concern; the integration of various research approaches to address this question, and estimate key quantities, is an on-going research challenge [101–103]. Risk to humans increases if there is any non-zero proportion of mutant viruses in the spillover inoculum, so tracking the presence of such mutants is beneficial. Surprisingly, the quantitative frequency of mutants in the initial dose has little impact on emergence probability in most scenarios, with the one exception of long-term infections with *α <* 1. Collectively, these results suggest that any knowledge of the cross-scale reproductive number *α* and mutant reproductive numbers can help to refine our goals for genetic surveillance, and that in many circumstances presence/absence detection is sufficient.

While there are not sufficient data from past emergence events to test our model’s conclusions, recent studies combining animal transmission experiments with deep sequencing have exhibited many phenomena aligned with our findings. Moncla et al. [43] conducted deep sequencing analyses of H1N1 influenza viruses, in the context of ferret airborne transmission experiments that examined the adaptation of avian-like viruses to the mammalian host. Their results provide in-depth insights into selection within hosts and at transmission bottlenecks, for a range of mutations on genetic backgrounds that change as adaptation proceeds (i.e. equivalent to numerous separate implementations of our model of a single mutational step). They observe a fascinating range of dynamics: some mutations appeared to have *α* moderately above 1, exhibiting modest increases in frequency between generations, but achieved this outcome with different traits (e.g. S113N on the HA190D225D background exhibited strong within-host selection and no evident transmission advantage, while D265V showed weak within-host selection but its frequency rises in transmission). Another mutation (I187T on the ‘Mut’ background) appeared to have *α* ≫ 1 and exhibited strong selection at both scales; notably, this mutation is widespread in 17/17 human-derived isolates of the post-emergence 1918 virus, consistent with the successful and rapid emergence our model would predict. Moncla et al. also present substantial evidence of cross-scale conflict in selection, as one mutation (G225D on ‘Mut’ background) exhibited declining frequencies within ferrets but rose to fixation in 2/2 transmission events, while numerous mutations in the HA2 region rose in frequency within the host but were eliminated in transmission. Another study examined a set of ‘gain-of-function’ mutations in H5N1 influenza in ferrets, and reported a slow rise in frequency when the virus was passaged between ferrets by intranasal inoculation, then rapid fixation of these mutations during airborne transmission [24]; the airborne transmission data are consistent with strong between-host selection and a high *α* value (though we emphasize that circulating H5N1 viruses required substantial modification to the favorable genetic background used in those experiments). Intriguingly, Moncla et al. synthesized their results with those of earlier studies [41, 49, 62] to hypothesize that the ‘stringency’ of the transmission bottleneck varies systematically during the course of viral adaptation, with loose bottlenecks prevailing when viruses first encounter a new host species (and perhaps again when the virus is host-adapted), and much tighter bottlenecks at the key juncture in host adaptation when a genotype with greater transmissibility is available to be selected. If this hypothesis is correct, then our findings can be applied to each adaptive step independently, and may help to identify which viral traits are most crucial to adaptive steps subject to tighter or looser bottlenecks.

Our results focus on systems where there is one major rate-limiting step to emergence, and the viral population can be represented by one wild-type and one mutant strain. This is a simplification of most viral emergence problems, but will apply directly to systems where a single large-effect mutation is the primary barrier to emergence of a supercritical strain, as for Venezuelan equine encephalitis virus emerging from rodents to horses [104]. While it is possible to extend our exact computations and analysis of the cross-scale reproductive number of a mutant virion to systems with multiple mutational steps, the present analysis already provides insights into more complex evolutionary scenarios. For evolutionary trajectories that proceed through a fixed series of genotypes, the probability of emergence can be approximated by extension of our equation (4), as in previous work [12, 13, 15]. If emergence requires multiple mutational steps which pass through a fitness valley, then the scale at which this valley occurs matters. A within-host fitness valley in replication rates would hinder pathogens with long-term infections and larger bottleneck widths, more than those with smaller bottlenecks. A between-host fitness valley in transmissibility could hinder evolutionary emergence of pathogens causing long-term infections more than those causing short-term infections, unless the within-host landscape is sufficiently favorable to allow traversing the valley within a single host’s long-term infection. Recent studies have also highlighted the importance of considering the broader genotype space, which can reveal indirect paths that circumvent fitness valleys [105], alternative genotypes that yield similar phenotypes [43], and the costs imposed by deleterious mutants on higher mutation rates [106].

Our analysis also focuses on a simple “logistic-like” model for within-host viral dynamics. This simplification allows us to study how evolutionary emergence is limited by different factors for pathogens dominated by exponential versus saturated phases of viral growth, while maintaining analytical tractability. Future important extensions would be to allow within-host fitness to alter the carrying capacity in the saturated phase, as well as identifying the relative contributions of stochastic within-host dynamics [17], immune responses, and host heterogeneity on viral emergence. Some of these aims would be addressed by using a more mechanistic model for the within-host dynamics, incorporating resource limitation [72, 73] or immune pressure [107]. We have assumed that the bottleneck width *N* is fixed for a given pathogen. This is broadly consistent with currently available data [44, 47, 48], but it will be important to explore the consequences of variation in bottleneck width arising from different routes of transmission, or possibly from changing viral loads [38, 39]. Among other possible impacts, this may alter the conclusion that emergence probability is minimally affected by the functional dependence of transmission probability on viral load. The computational and analytical framework developed here can be extended to account for these additional complexities. Other important extensions can explore the impact of clonal competition on emergence probabilities [108–111] or the potential for complementation to rescue pathogen strains from deep fitness valleys–a mechanism that depends on wide transmission bottlenecks [112].

Our cross-scale analysis opens the door for a new generation of integrative risk assessment models for pathogen emergence, which will integrate growing streams of data collected in laboratories and field surveillance programs [100, 101, 103]. At present we rely on the intuition of individual scientists to link together the discoveries from targeted experiments, massively parallel phenotypic screens, experimental evolution, clinical medicine, and field epidemiology and disease ecology. Mathematical and computational models that connect biological scales using mechanistic principles can make unique contributions to this transdisciplinary enterprise, by formally integrating diverse empirical findings and by identifying the crucial knowledge gaps to focus future research. The work presented here is a step on the path to realizing this potential.

## Acknowledgments

The authors thank two anonymous reviewers and Joshua Weitz for providing valuable feedback that greatly improved the quality of this paper. This work was supported by U.S. National Science Foundation Grants EF-0928987 and DMS-1716803 to SJS, and EF-0928690 and DEB-1557022 to JLS, and DARPA PREEMPT D18AC00031 to JLS. The content of the information does not necessarily reflect the position or the policy of the U.S. government, and no official endorsement should be inferred.

## Appendix

### Derivation of the Single Strain Reproductive Numbers

When the within-host dynamics only exhibit exponential growth (i.e. *N* exp(*r*_*i*_*T*) < *K*) and there is a linear transmission function, the basic reproductive numbers equal

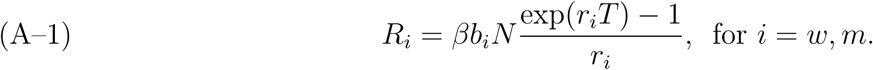

When the within-host dynamics saturate (i.e. *N* exp(*r*_*i*_*T*) > *K*), the basic reproductive number equals

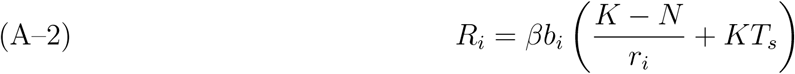

where *T*_*e*_ = log(*K/N*)*/r*_*i*_ is the length of exponential phase and *T*_*s*_ = *T* − *T*_*e*_ is the length of saturated phase. These expressions for the single strain reproductive numbers are equivalent to evaluating the integral presented in equation (22) of Coombs et al. [73] for our within-host model. We also note that our assumption of small mutation rates and *R*_*w*_ < *R*_*m*_ implies that the multiscale reproductive number *R*_0_ in the sense of Coombs et al. [73] and Lythgoe et al. [65] (i.e. the dominant eigenvalue of the next generation matrix of *DG*(1, 1, …, 1) is (approximately) *R*_*m*_.

We derive two approximations of *R*_*i*_ under the assumption that *s* is small, exp(*r*_*i*_*T*) ≫ 1, and *K* ≫ *N*. First, assume that the infection is short-term in which case *T*_*e*_ = *T*. Then provided *s* is sufficiently small to ensure that the mutant type doesn’t saturate, *R*_*i*_ are given by (A–1). The log ratio, provided exp(*r*_*i*_*T*) ≫ 1, satisfies

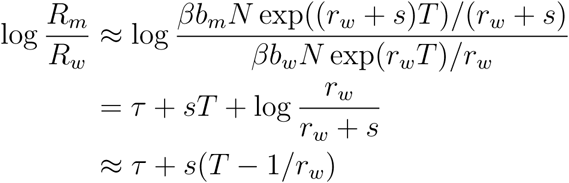

which yields (2) in the main text.

Now assume that the infection is long-term in which case *T*_*e*_ < *T*, and that if *s* < 0, |*s*| is sufficiently small to ensure that the mutant type also saturates before time *T*. Then the basic reproductive numbers *R*_*i*_ are given by (A–2). If *K* ≪ *N*, then

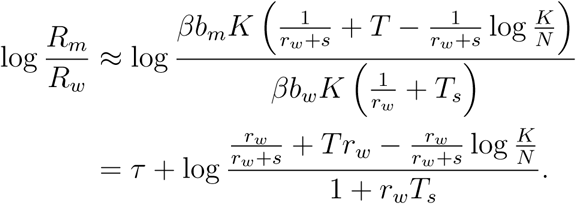

Assume that |*s*| ≪ *r*_*w*_. Then

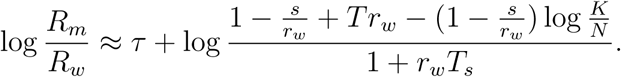

As *T* = *T*_*s*_ + *T*_*e*_ and

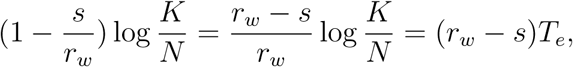

it follows that

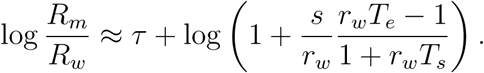

As log(1 + *x*) ≈ *x* for small *x* and |*s*| ≪ *r*_*w*_ by assumption, we obtain

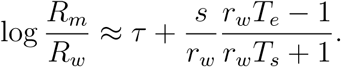

Equation (3) in the main text follows in the case that *T*_*s*_ ≫ *T*_*s*_ in which case the second term is approximately zero.

### Derivation of the Emergence Probability Approximation

For small mutation likelihood *μ*, we derive a mathematically explicit version of the approximation (4) for the emergence probability from the main text. As stated in the main text, this approximation is given by the product of three terms: the expected number of secondary, wild-type cases produced during a fade-out, the mean number of individuals infected with mutant virions by an individual initially infected only with the wild-type, and the probability of emergence from an individual infected with a single mutant virion. As noted in the main text, the first term is given by 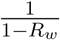. The second term requires more work. To derive an analytic approximation for this term, notice that the mean number of individuals infected with *ℓ* mutant virions by an individual only infected with the wild type equals

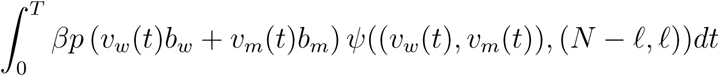

where *v*_*w*_(*t*), *v*_*m*_(*t*) is the solution of the within host viral dynamics with *v*_*w*_(0) = *N, v*_*m*_(0) = 0, and 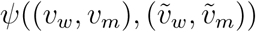 is the probability of an individual with viral load (*v*_*w*_, *v*_*m*_) infecting an individual with a viral load of 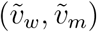 where 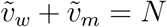. The solution (*v*_*m*_(*t*), *v*_*m*_(*t*)) is given by

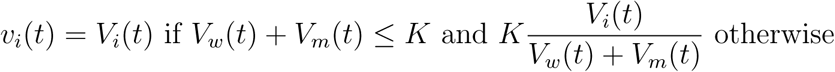

where *V*_*w*_(*t*), *V*_*m*_(*t*) are the solutions to

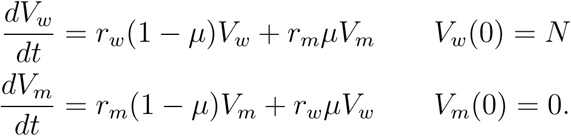

Ignoring back mutations (i.e. setting *r*_*m*_*μ* = 0 and *r*_*m*_(1 − *μ*) to *r*_*m*_), the solutions for *V*_*w*_(*t*), *V*_*m*_(*t*) are approximately

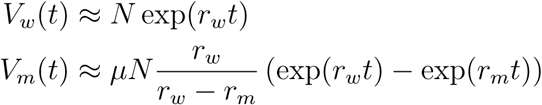

if *r*_*w*_ ≠ *r*_*m*_, and

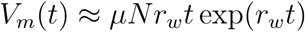

if *r*_*w*_ = *r*_*m*_ = *r*. Since 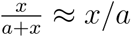 to first order near 0, the weighted frequency, *x*_*m*_(*t*), of mutant strain is approximately

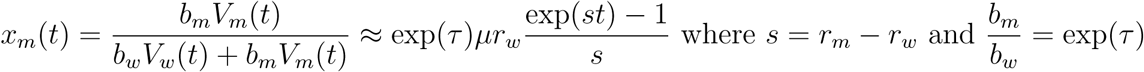

if *s* ≠ 0, and

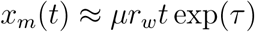

if *s* = 0. We have

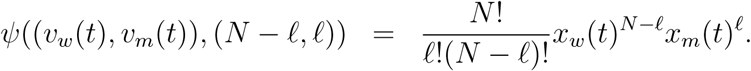

For *ℓ* ≥ 2, these terms are of order *μ*^2^ and therefore will be ignored. Hence, the only term of interest is *ℓ* = 1:

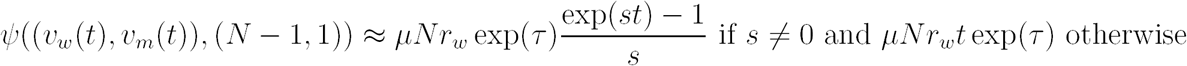

We also can approximate (assuming *p* is differentiable)

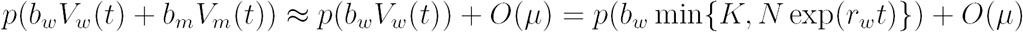

We drop the *O*(*μ*) term as it will only lead to higher order terms in the approximation.

Putting all of this together gives the following approximation for the mutant force of infection

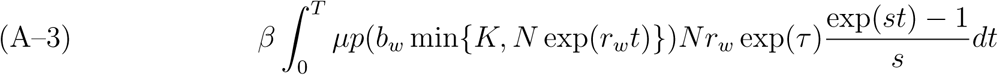

if *s* ≠ 0, and

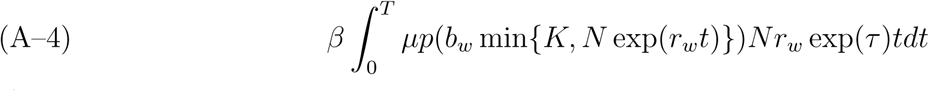

if *s* = 0.

In the case of a linear transmission function *p*(*x*) = *x*, we can write down explicit expressions for (A–3) and (A–4). There are two cases to consider. First suppose that *N* exp(*r*_*w*_*T*) ≤ *K* i.e. the infection is short-term. Then, integrating and simplifying yields the following approximation for the mutant force of infection

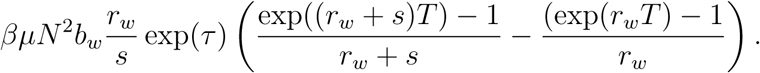

Assuming *r*_*w*_ ≫ *s* (and thus *r /*(*r*_*w*_ + *s*) ≃ 1 − *s/r*_*w*_),

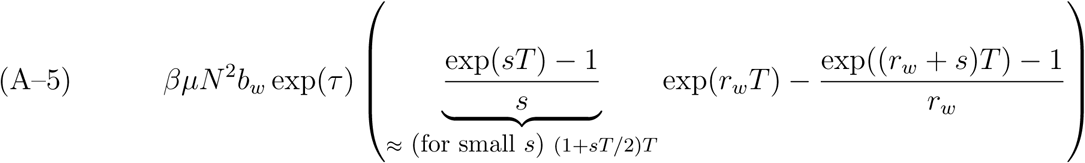

if *s* ≠ 0 and

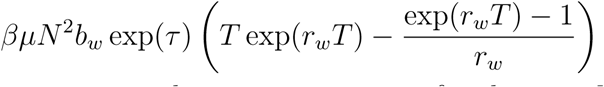

if *s* = 0. Now rather than writing out the entire expression for the case *N* exp(*r*_*w*_*T*) ≥ *K*, lets write down things for when the time in the saturated phase is much, much longer than the time in the exponential phase. Then, integrating and simplifying yields the following approximation for the mutant force of infection

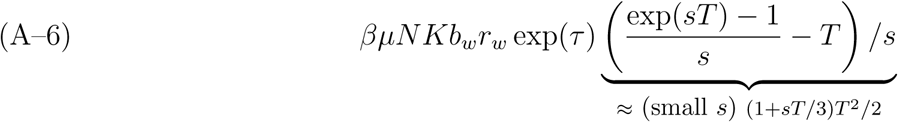

if *s* ≠ 0, and

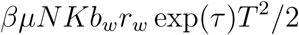

otherwise.

Putting this all together, (4) for an short-term (respectively long-term) infection with *s* ≠ 0 becomes the product of 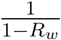, (A–5) (respectively (A–6)), and the probability of an outbreak starting with one individual infected with *N* − 1 wild type virions and 1 mutant type virions. The final probability term can be calculated exactly using the generating functions described in the <u>Models and Methods</u> section of the main text. Fig Appendix–6 illustrates the effectiveness of this approximation, and Fig Appendix–7 plots the the error in the approximation.

### Derivation of the Mean Field Frequency Dynamics

To understand how the viral composition of infected individuals change across generations, we derive a mean field approximation for the dynamics of the mean mutant viral load at the beginning of each generation of disease spread. To this end, we define a map from *h* : [0, 1] → [0, 1] where *x* ∈ [0, 1] represents the current mean mutant viral load in the population at the beginning of the infectious period and *h*(*x*) is the mean at the beginning of infectious period in the next generation. Our derivation of this mean field dynamic is done in the limit of large *N* ↑ ∞ and *μ* ↓ 0. None-the-less, as shown by the dashed red line in Fig 2D, this approximation works quite well away from this limit.

We begin by approximating the mean initial mutant viral count in individuals infected by an individual with *V*_*w*_(0) = *N* − *ℓ* and *V*_*m*_(0) = *ℓ*. Recall, the force of infection for producing individuals initially infected with *j* mutant viral particles is given by

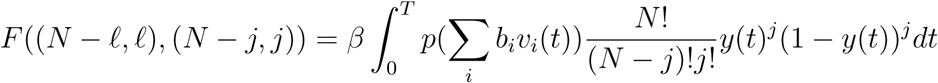

where 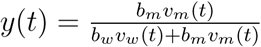 is the within-host frequency of the mutant strain, and (*v*_*w*_(*t*), *v*_*m*_(*t*)) is the solution of the within-host viral dynamics with initial condition *v*_*w*_(0) = *N* − *ℓ, v*_*m*_(0) = *ℓ*. Weighting this term by *j* and summing over *j* yields the expected number of mutant viral particles in an individual infected by our type (*N* − *ℓ, ℓ*) infected individual:

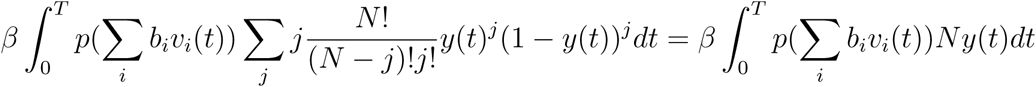

Now if we let *x* = *ℓ/N* denote the initial fraction, then dividing the previous integral by the net number of viral particles infecting new individuals yields our desired update rule

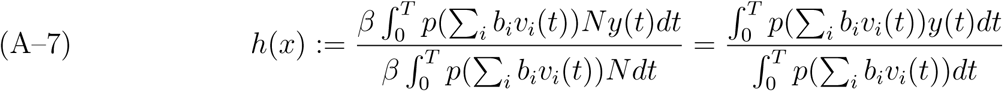

Note that *h*(*x*) is a function of *x* as the solution of (*V*_*w*_(*t*), *V*_*m*_(*t*)) depends on its initial condition *V*_*w*_(0) = (1 − *x*)*N, V*_*m*_(0) = *xN*.

The points *x* = 0 and *x* = 1 are fixed points for *h* corresponding to a mutant-free and wild-type-free states. Stability of the fixed point *x* = 0 is determined by

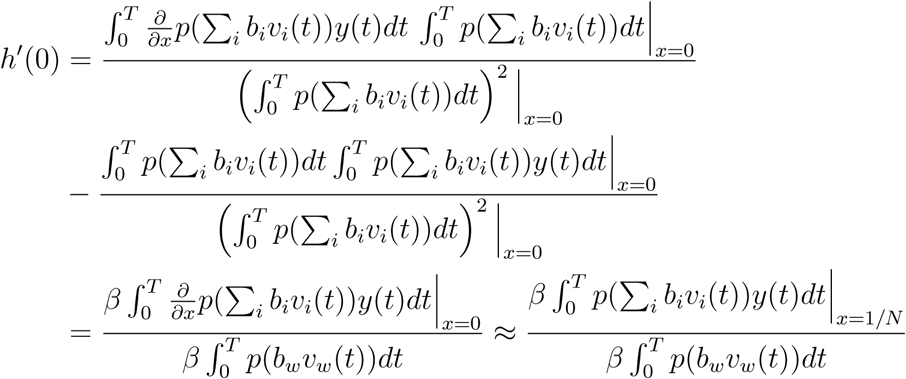

for *N* ≫ 1. *h*′(0) corresponds to *α* described in the main text and the final expression has the verbal interpretation given in the main text.

In the special case of a linear transmission function, *p*(*x*) = *x*, we get the simplified expression

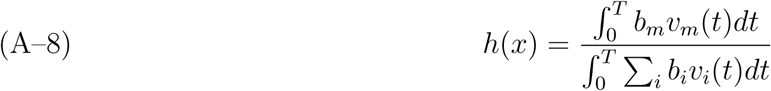

where

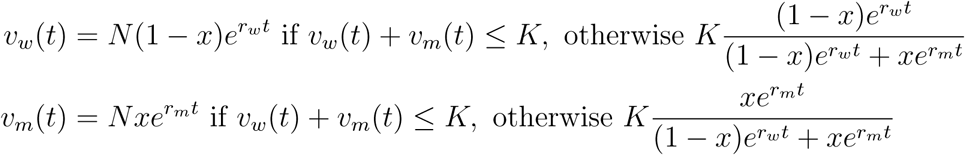

Carrying out the integration, in general, is complicated by the fact that the time at which *V*_*w*_(*t*) + *V*_*m*_(*t*) = *K* has no explicit formula when *s* ≠ 0 and, in general, this saturation time will depend on *x*.

In the special case of short-term infections (i.e. there is only exponential growth), we get

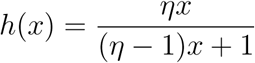

where

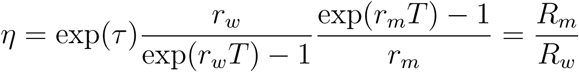

*α* is defined as *h*′ (0), which here is equal to *η*, thus for short-term infections, *α* = *R*_*m*_*/R*_*w*_.

Since *R*_*w*_ < 1 by assumption and *R*_*m*_ > 1 is necessary for emergence, we always have *α* > 1 and so the frequency dependent dynamics at the scale of the host population can not significantly impede emergence.

Now, lets consider the more difficult case of a long-term infection with a saturated phase to the within-host viral dynamics. Then

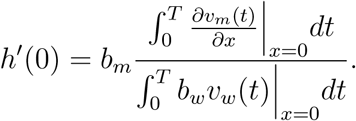

For *x* close to 0, we have the time at which the dynamics saturate, *T*_*e*_, is given approximately by

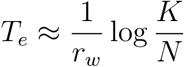

in which case

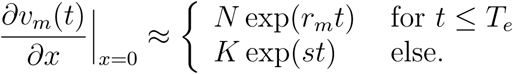

Let *T*_*s*_ = *T* −*T*_*e*_ and assume that 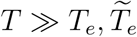 where 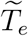 is the length of the exponential phase for an individual infected only with the mutant strain. Then

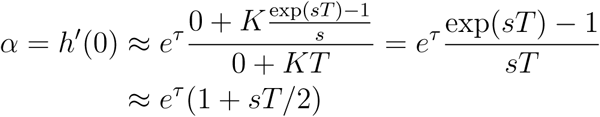

as claimed in the main text.

### Estimating the Probability of Emergence when

*α* < 1. When *α* is less than 1, the frequency of mutant virus decrease in an infected host, and consequently, even if the adapted virus may emerge, the probability of emergence is very low, and even lower when the bottleneck size, *N*, increases. Here, we provide an approximation for the emergence probability when *α* < 1, which explains why the probability of emergence decreases dramatically when *N* increases.

When the outbreak starts, the first individual is infected with wild-type only. When *s* < 0, the mutation-selection balance can be reached relatively quickly, and for *s* negative enough, the proportion of mutant is small. So the probability to transmit at least one mutant is roughly equal to the probability to transmit one mutant, which is *N* exp(*τ*)*r*_*w*_*μ/* |*s*| where *r*_*w*_*μ/* |*s*| is the proportion of the mutant type, and exp(*τ*) is its relative transmissibility. Then, if *s* is small enough, then the reproductive number of an individual with a mixed transmission is close to *R*_*w*_ of the wild-type. Thus, the number of transmissions in a wild-type outbreak can be used (*R*_*w*_*/*(1 − *R*_*w*_)). For an individual infected with a mixed infection, what will lead to emergence are the contacts for which only the mutant is transmitted. The number of such contacts is:

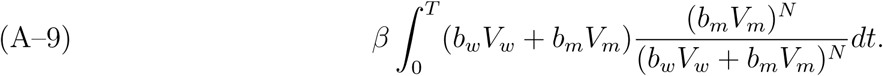

This can be re-written as:

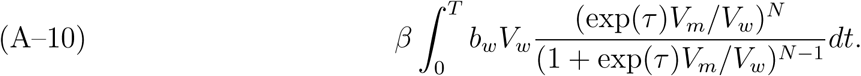

As most cases of mixed infection will be cases started with a mix of one mutant and *N* − 1 wild-type viral particles, *V*_*m*_*/V*_*w*_ = exp(*st*)*/*(*N* − 1). Thus previous expression is equal to:

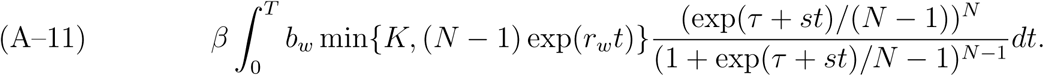

**Figure Appendix-1.**
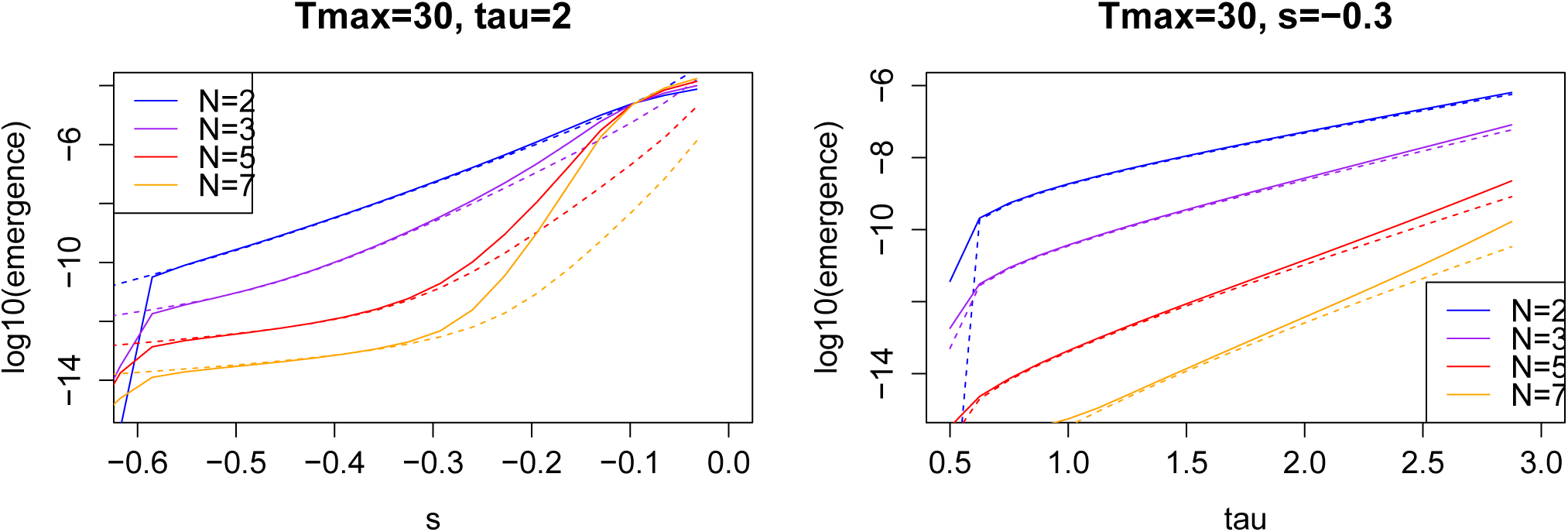
Approximation (A–12) (dashed lines) vs. numerical resolution of the generating maps (R code) (solid lines)

Last, an individual infected with mutant viruses alone has to lead to a successful outbreak, which happens at approximately the same probability than in the case with no back mutations, with probability *p*_*m*_. So overall, the approximation will be:

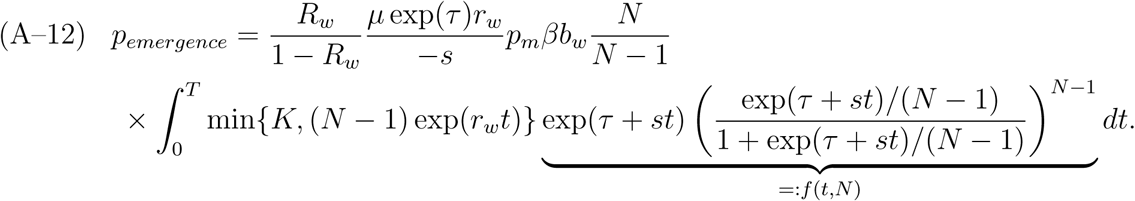

Now we can ask, which parts of this expression depend on *N*? The mutant reproductive number 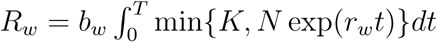 is independent from *N*, because we have chosen *bw* to keep *R*_*w*_ the same for all *N* values. Thus 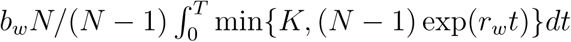 is almost independent from *N*. Therefore, most of the dependence of 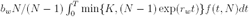 with *N* stems from the dependence of *f* (*t, N*) with *N*. Since *a* ↦ *a/*(1 + *a*) is an increasing function bounded above by 1 for positive *a*, the expression

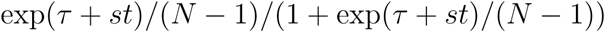

decreases when *N* increases. As *N* ↦ (*a/*(1 + *a*))^*N*− 1^ is a decreasing function of *N* ≥ 1 for *a* > 0, we get that the probability of emergence decreases at least exponentially with the bottleneck size, as claimed in the main text. Fig Appendix-1 illustrates that these approximations work especially when *s* is sufficiently negative.

#### Numerics with Nonlinear Transmission Functions

To explore the robustness of our numerical results to the assumption of a linear transmission function, we redid our numerical analysis with two non-linear transmission functions *p*(*x*) = 1 − exp (− *x*) in Fig Appendix–2 and *p*(*x*) = log(1 + *x*) in Fig Appendix–4. Differences between the emergence probabilities for the nonlinear and linear transmission functions are shown in Figs Appendix–3 through 4. As these figures demonstrate, we nearly get the same results.

**Figure Appendix–2.**
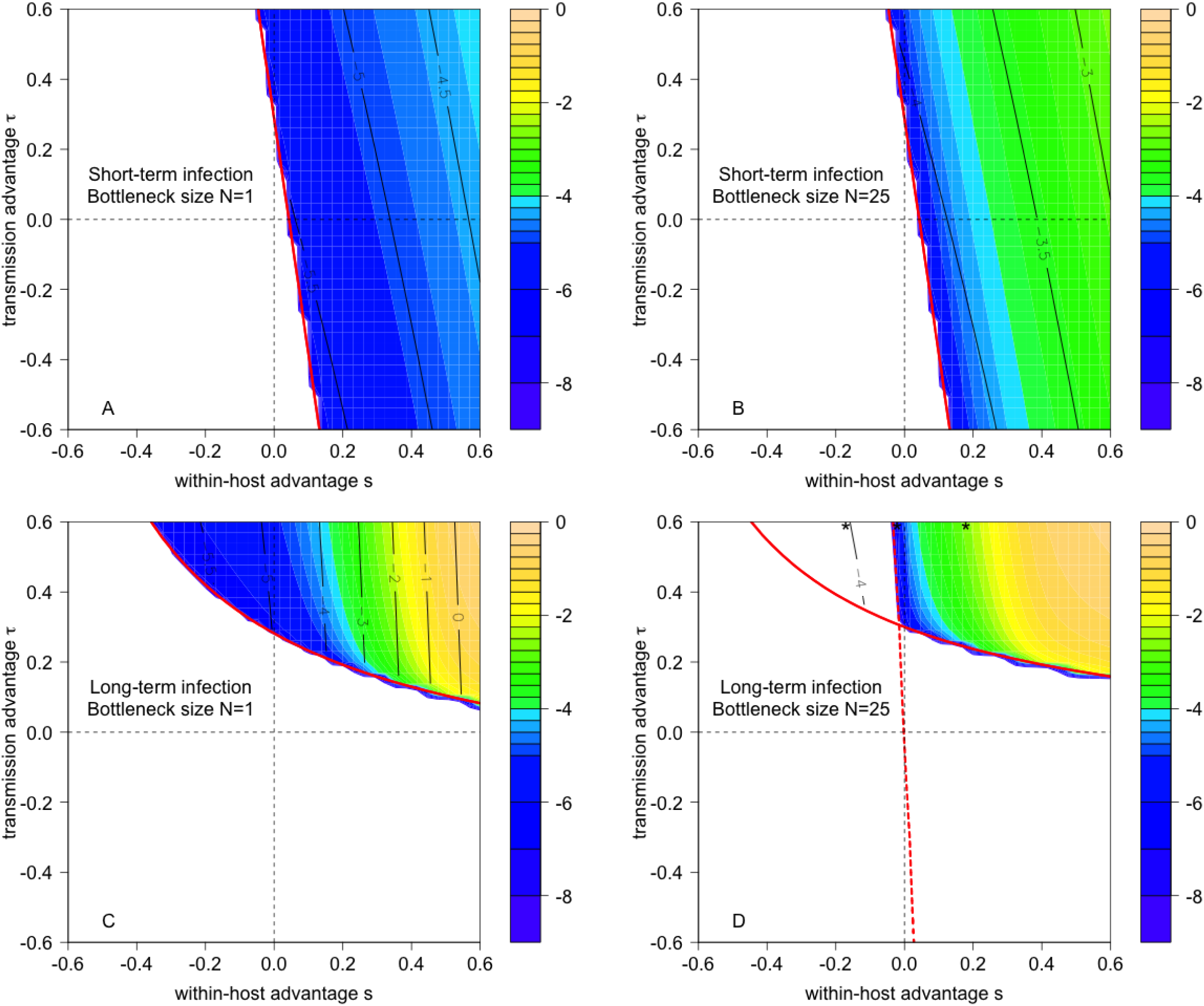
Emergence probabilities for the transmission function *p*(*x*) = 1 − exp(− *x*) with all other parameters as indicated in Fig 2.

**Figure Appendix–3.**
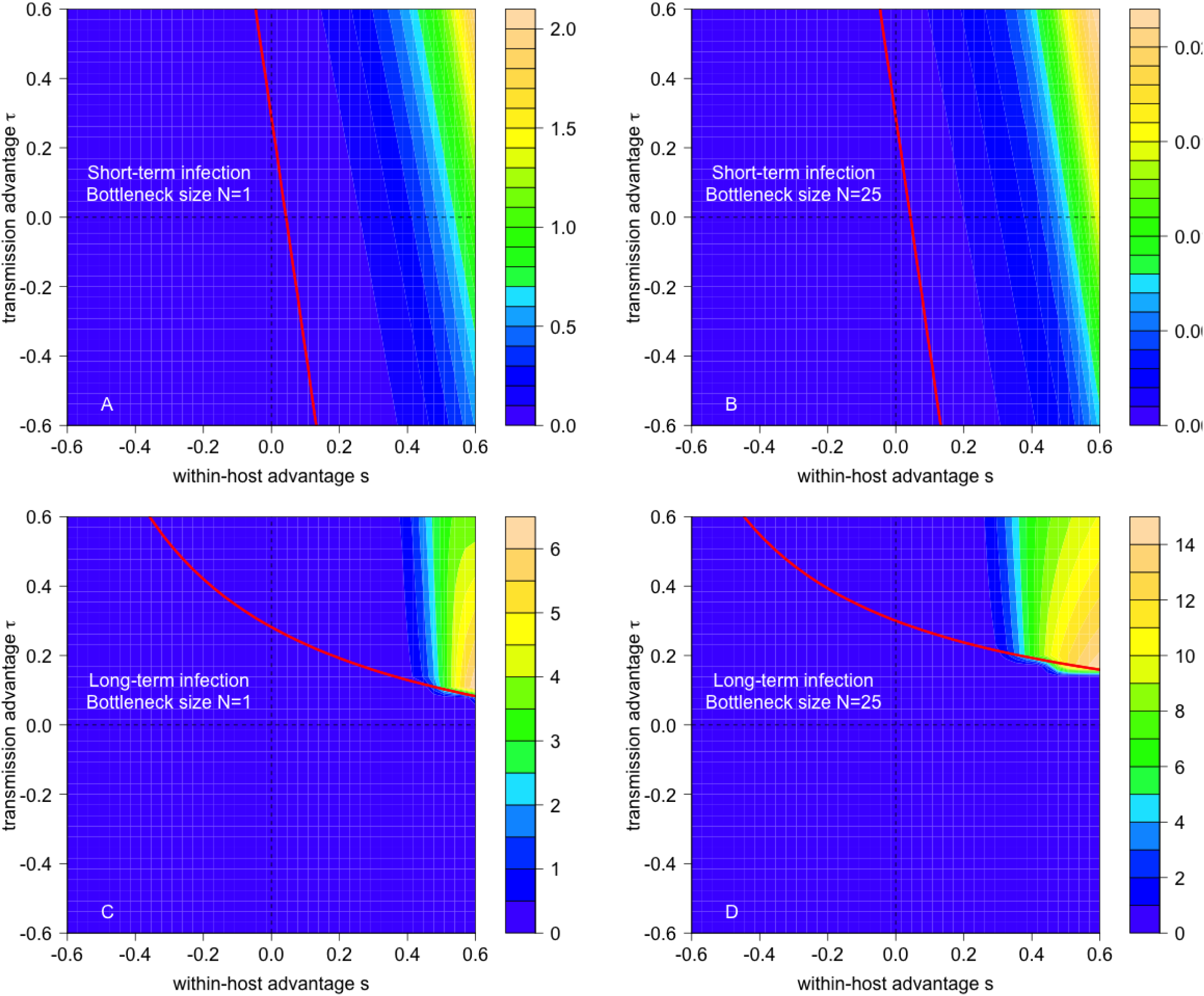
Contour plots of 1, 000 × the absolute value of the difference between the emergence probabilities for the transmission functions *p*(*x*) = 1 −exp (−*x*) and *p*(*x*) = *x*. Parameters as indicated in Fig 2.

**Figure Appendix–4.**
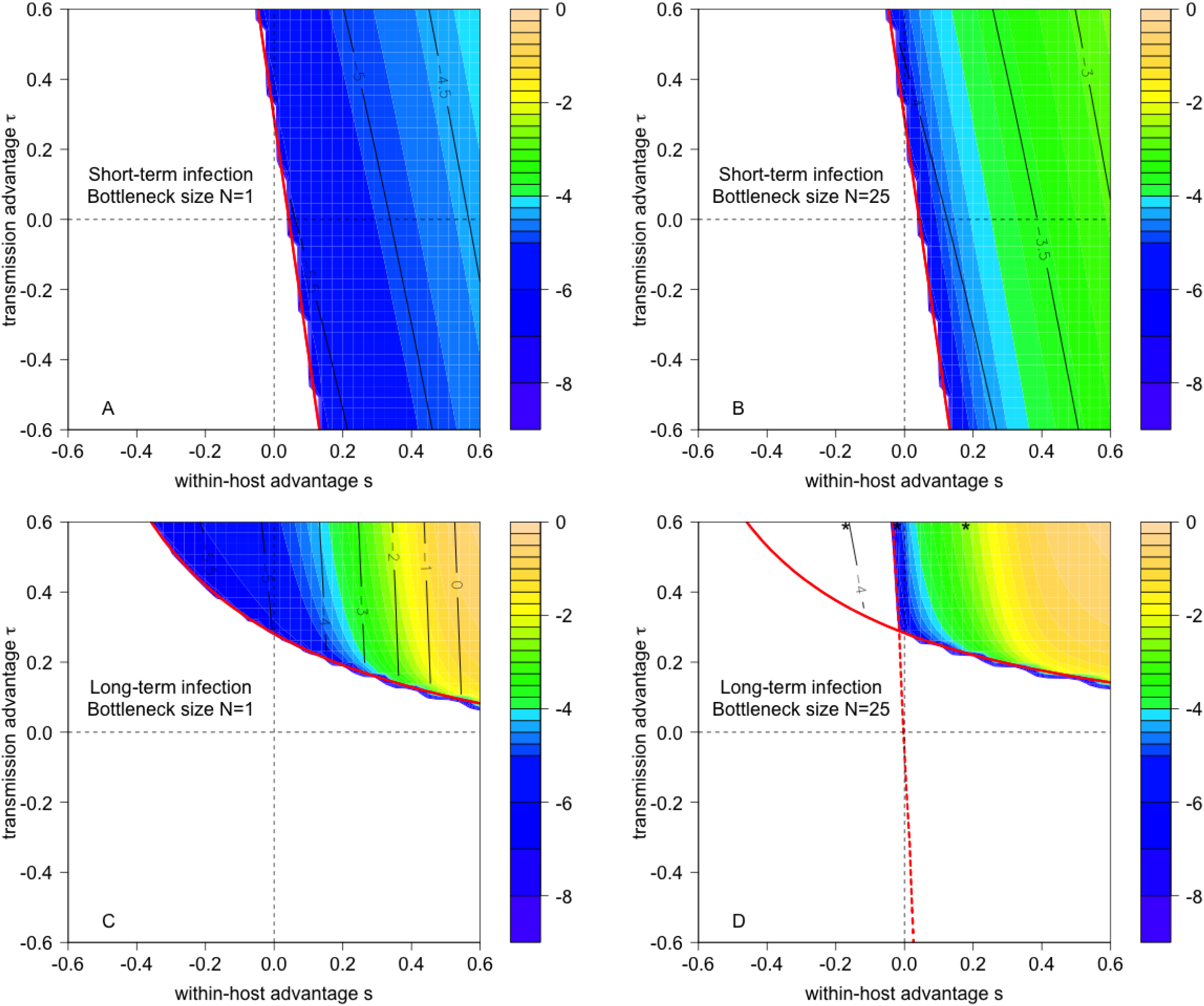
Emergence probabilities for the transmission function *p*(*x*) = log(1 + *x*) with all other parameters as indicated in Fig 2.

**Figure Appendix–5.**
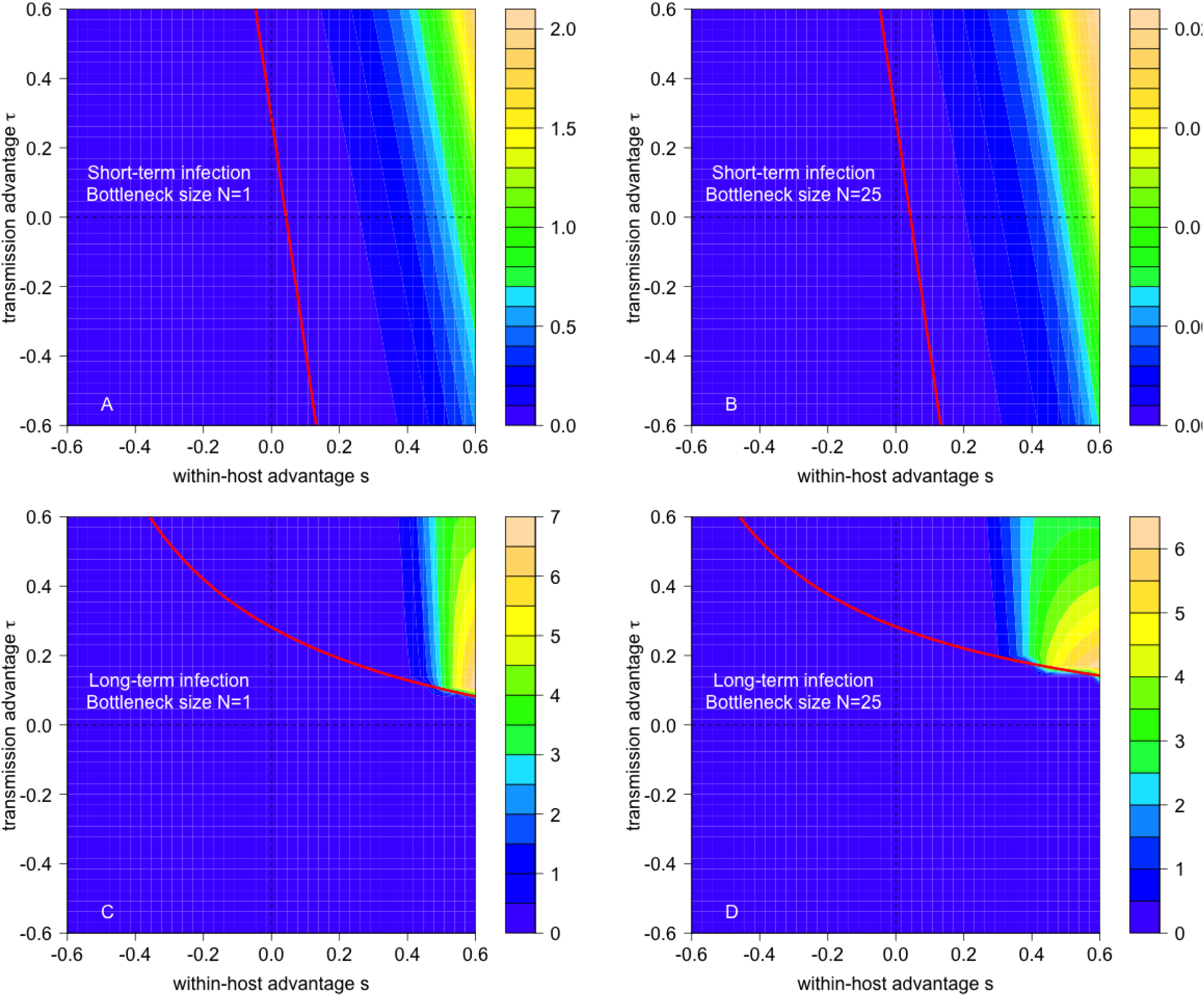
Contour plots of 1, 000 × the absolute value of the difference between the emergence probabilities for the transmission functions *p*(*x*) = log(1 + *x*) and *p*(*x*) = *x*. Parameters as indicated in Fig 2.

**Figure Appendix–6.**
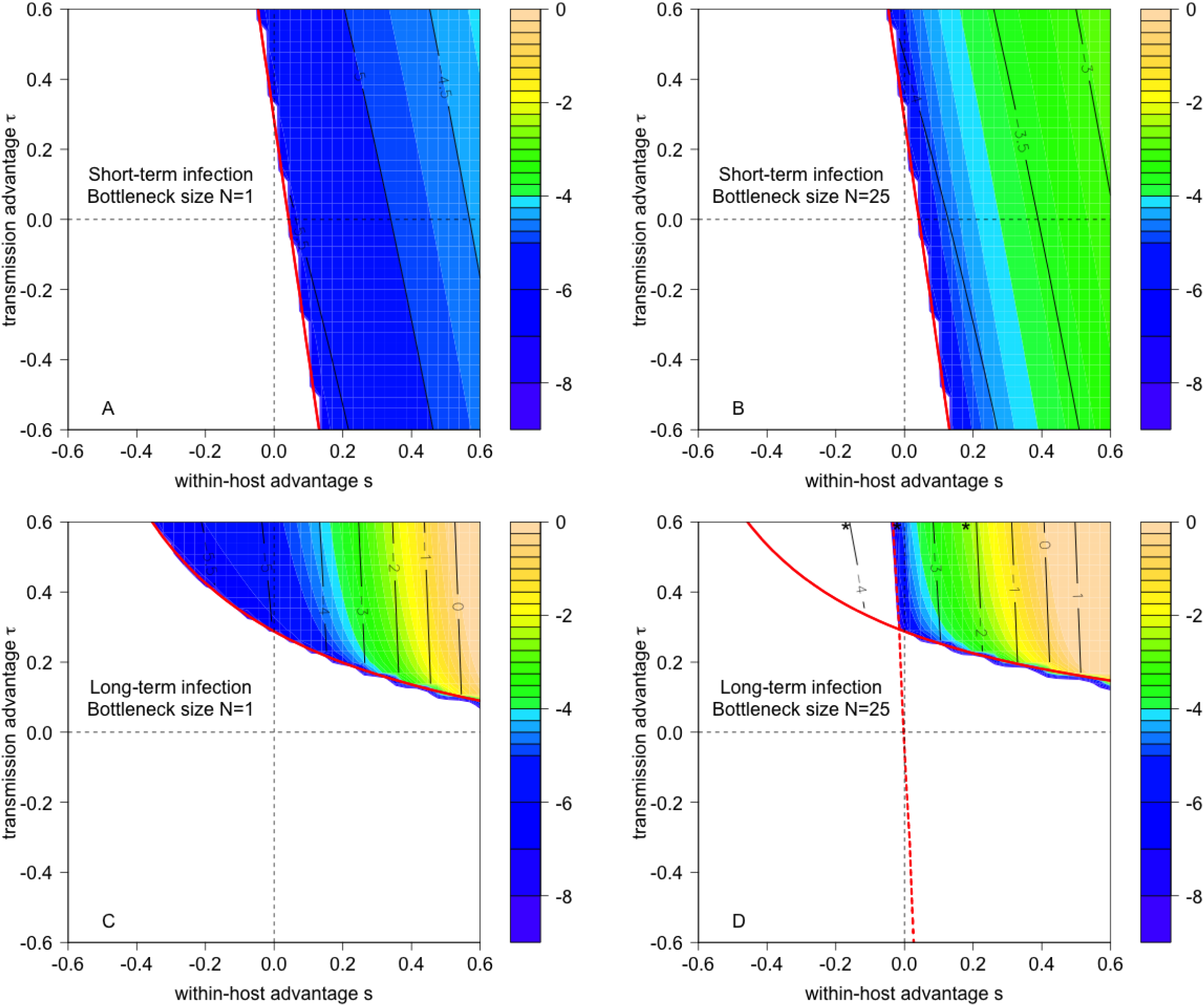
The analytic approximation based on (4) for the exact computations of the emergence probabilities in Fig 2 in the main text.

**Figure Appendix–7.**
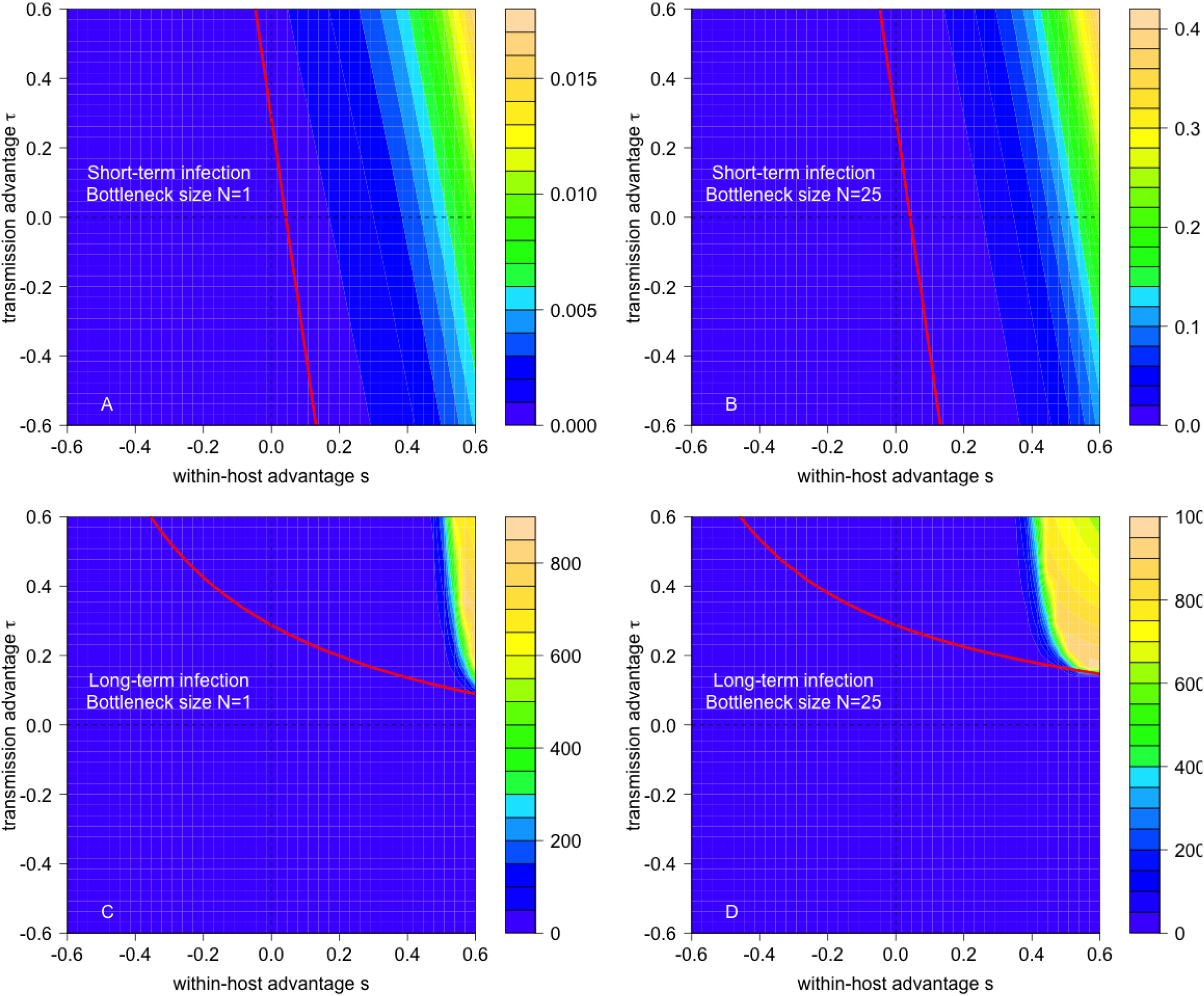
Contour plots of 1, 000 × the absolute value in the difference between the analytic approximation from (4) and the exact computations for the emergence probabilities. Parameters as in Fig 2 in the main text.

## References

[1] K.E. Jones, N.G. Patel, M.A. Levy, A. Storeygard, D. Balk, J.L. Gittleman, and P. Daszak. Global trends in emerging infectious diseases. Nature, 451:990–993, 2008.

[2] M. Woolhouse, F. Scott, Z. Hudson, R. Howey, and M. Chase-Topping. Human viruses: discovery and emergence. Philosophical Transactions of the Royal Society B: Biological Sciences, 367:2864–2871, 2012.

[3] S.S. Morse, J.A.K. Mazet, M. Woolhouse, C.R. Parrish, D. Carroll, W.B. Karesh, C. Zambrana-Torrelio, W.I. Lipkin, and P. Daszak. Prediction and prevention of the next pandemic zoonosis. The Lancet, 380:1956–1965, 2012.

[4] C.R. Howard and N.F. Fletcher. Emerging virus diseases: can we ever expect the unexpected? Emerging Microbes and Infections, 1:e46, 2012.

[5] J. Steel, A.C. Lowen, S. Mubareka, and Palese P. Transmission of influenza virus in a mammalian host is increased by pb2 amino acids 627k or 627e/701n. PLOS Pathogens, 5: e1000252, 2009.

[6] B.V. Lowder, C.M. Guinane, N. L. Ben Zakour, L.A. Weinert, A. Conway-Morris, R.A. Cartwright, A.J. Simpson, A. Rambaut, U. Nübel, and J.R. Fitzgerald. Recent human-to-poultry host jump, adaptation, and pandemic spread of Staphylococcus aureus. Proceedings of the National Academy of Sciences, 106:19545–19550, 2009.

[7] L. Xu, L. Bao, W. Deng, H. Zhu, F. Li, T. Chen, Q. Lv, J. Yuan, Y. Xu, Y. Li, Y. Yao, S. Gu, P. Yu, H. Chen, and C. Qin. Rapid adaptation of avian h7n9 virus in pigs. Virology, 452-453:231–236, 2014.

[8] M. Jonges, M.R.A. Welkers, R.E. Jeeninga, A. Meijer, P. Schneeberger, R.A.M. Fouchier, M.D. de Jong, and M. Koopmans. Emergence of the virulence-associated pb2 e627k substitution in a fatal human case of highly pathogenic avian influenza virus a (h7n7) infection as determined by illumina ultra-deep sequencing. Journal of virology, 88(3): 1694–1702, 2014. ISSN 0022-538X.

[9] W. Zhu and Y. Shu. Genetic tuning of avian influenza A (H7N9) virus promotes viral fitness within different species. Microbes and Infection, 17:118–122, 2015.

[10] T. Lam, B. Zhou, J. Wang, Y. Chai, Y. Shen, X. Chen, C. Ma, W. Hong, Y. Chen, Y. Zhang, L. Duan, P. Chen, J. Jiang, Yu Zhang, Lifeng Li, Leo Lit Man Poon, Richard J. Webby, David K. Smith, Gabriel M. Leung, Joseph S. M. Peiris, Edward C. Holmes, Yi Guan, and Huachen Zhu. Dissemination, divergence and establishment of H7N9 influenza viruses in china. Nature, 522:102–105, 2015.

[11] Dan Xiang, Xuejuan Shen, Zhiqing Pu, David M Irwin, Ming Liao, and Yongyi Shen. Convergent evolution of human-isolated h7n9 avian influenza a viruses. The Journal of infectious diseases, 217(11):1699–1707, 2018.

[12] R. Antia, R.R. Regoes, J.C. Koella, and C.T. Bergstrom. The role of evolution in the emergence of infectious diseases. Nature, 426:658–661, 2003.

[13] Y. Iwasa, F. Michor, and M.A. Nowak. Evolutionary dynamics of invasion and escape. Journal of Theoretical Biology, 226:205–214, 2004.

[14] J.B. André and T. Day. The effect of disease life history on the evolutionary emergence of novel pathogens. Proceedings of the Royal Society B: Biological Sciences, 272:1949–1956, 2005.

[15] M. Park, C. Loverdo, S.J. Schreiber, and J.O. Lloyd-Smith. Multiple scales of selection influence the evolutionary emergence of novel pathogens. Philosophical Transactions of the Royal Society B: Biological Sciences, 368:20120333, 2013.

[16] K.M. Peck, C.H.S. Chan, and M.M. Tanaka. Connecting within-host dynamics to the rate of viral molecular evolution. Virus Evolution, 1:vev013, 2015.

[17] J.L. Geoghegan, A.M. Senior, and E.C. Holmes. Pathogen population bottlenecks and adaptive landscapes: overcoming the barriers to disease emergence. Proceedings of the Royal Society B, 283:20160727, 2016.

[18] B. T. Grenfell, O. G. Pybus, J. R. Gog, J. L. N. Wood, J. M. Daly, J. A. Mumford, and E. C. Holmes. Unifying the epidemiological and evolutionary dynamics of pathogens. Science, 303:327–332, 2004.

[19] O. G. Pybus and A. Rambaut. Evolutionary analysis of the dynamics of viral infectious disease. Nature Reviews Genetics, 10:540–550, 2009.

[20] P. S. Pennings, S. Kryazhimskiy, and J. Wakeley. Loss and recovery of genetic diversity in adapting populations of HIV. PLOS Genetics, 10:e1004000, 2014.

[21] S. G. Lim, Y. Cheng, S. Guindon, B. L. Seet, L. Y. Lee, P. Hu, S. Wasser, F. J. Peter, T. Tan, and M. Goode. Viral quasi-species evolution during hepatitis Be antigen seroconversion. Gastroenterology, 133:951–958, 2007.

[22] Gustavo H Kijak, Eric Sanders-Buell, Agnes-Laurence Chenine, Michael A Eller, Nilu Goonetilleke, Rasmi Thomas, Sivan Leviyang, Elizabeth A Harbolick, Meera Bose, Phuc Pham, et al. Rare hiv-1 transmitted/founder lineages identified by deep viral sequencing contribute to rapid shifts in dominant quasispecies during acute and early infection. PLoS pathogens, 13(7):e1006510, 2017.

[23] E. Domingo, J. Sheldon, and C. Perales. Viral quasispecies evolution. Microbiology and Molecular Biology Reviews, 76:159–216, 2012.

[24] M. Linster, S. van Boheemen, M. de Graaf, E.J.A. Schrauwen, P. Lexmond, B. Mänz, T.M. Bestebroer, J. Baumann, D. van Riel, and G.F. Rimmelzwaan. Identification, characterization, and natural selection of mutations driving airborne transmission of a/h5n1 virus. Cell, 157:329–339, 2014.

[25] P. Lemey, A. Rambaut, and O. G. Pybus. Hiv evolutionary dynamics within and among hosts. AIDS Reviews, 8:125–140, 2006.

[26] E. C. Holmes. Viral evolution in the genomic age. PLOS Biology, 5:2104–2105, 2007.

[27] P. R. Murcia, G. J. Baillie, J. Daly, D. Elton, C. Jervis, J. A. Mumford, R. Newton, C. R. Parrish, K. Hoelzer, and G. Dougan. Intra-and interhost evolutionary dynamics of equine influenza virus. Journal of Virology, 84:6943–6954, 2010.

[28] R. A. Bull, J. S. Eden, F. Luciani, K. McElroy, W. D. Rawlinson, and P. A. White. Contribution of intra-and interhost dynamics to norovirus evolution. Journal of Virology, 86:3219–3229, 2012.

[29] J. Hughes, R. C. Allen, M. Baguelin, K. Hampson, G. J. Baillie, D. Elton, J. R. Newton, P. Kellam, J. L. Wood, E. C. Holmes, and P. R. Murcia. Transmission of equine influenza virus during an outbreak is characterized by frequent mixed infections and loose transmission bottlenecks. PLOS Pathogens, 8:e1003081, 2012.

[30] J.C. Stack, P.R. Murcia, B.T. Grenfell, J.L. Wood, and E.C. Holmes. Inferring the inter-host transmission of influenza a virus using patterns of intra-host genetic variation. Proceedings of the Royal Society B: Biological Sciences, 2012.

[31] M. J. Morelli, C. F. Wright, N. J. Knowles, N. Juleff, D. J. Paton, D. P. King, and D. T. Haydon. Evolution of foot-and-mouth disease virus intra-sample sequence diversity during serial transmission in bovine hosts. Veterinary Research, 44:1–15, 2013.

[32] R. J. Orton, C. F. Wright, M. J. Morelli, N. Juleff, G. Theibaud, N. J. Knowles, B. Valdazo-Gonzalez, D. J. Paton, D. P. King, and D. T. Haydon. Observing micro-evolutionary processes of viral populations at multiple scales. Philosophical Transactions of the Royal Society B: Biological Sciences, 368, 2013.

[33] B. Vrancken, A. Rambaut, M. A. Suchard, A. Drummond, G. Baele, I. Derdelinckx, Van Wijngaerden, Vandamme E., A. M., Van Laethem, Lemey K., and P. The Genealogical Population Dynamics of HIV-1 in a Large Transmission Chain: Bridging within and among Host Evolutionary Rates. PLOS Computational Biology, 10:e1003505, 2014.

[34] Bram Vrancken, Marc A Suchard, and Philippe Lemey. Accurate quantification of within- and between-host HBV evolutionary rates requires explicit transmission chain modelling. Virus Evolution, 3(2), 10 2017. ISSN 2057-1577. doi: 10.1093/ve/vex028. URL https://doi.org/10.1093/ve/vex028.vex028.

[35] E. Duarte, D. Clarke, A. Moya, E. Domingo, and J. Holland. Rapid fitness losses in mammalian RNA virus clones due to Muller’s ratchet. Proceedings of the National Academy of Sciences, 89:6015–6019, 1992.

[36] C. T. Bergstrom, P. McElhany, and L. A. Real. Transmission bottlenecks as determinants of virulence in rapidly evolving pathogens. Proceedings of the National Academy of Sciences, 96:5095–5100, 1999.

[37] I. S. Novella, J. Quer, E. Domingo, and J. J. Holland. Exponential fitness gains of RNA virus populations are limited by bottleneck effects. Journal of Virology, 73:1668–1671, 1999.

[38] Mark P Zwart and Santiago F Elena. Matters of size: genetic bottlenecks in virus infection and their potential impact on evolution. Annual Review of Virology, 2:161–179, 2015.

[39] John T McCrone and Adam S Lauring. Genetic bottlenecks in intraspecies virus transmission. Current opinion in virology, 28:20–25, 2018.

[40] J.S. LeClair and L.M. Wahl. The impact of population bottlenecks on microbial adaptation. Journal of Statistical Physics, 172:114–125, 2018.

[41] A. Varble, R. A. Albrecht, S. Backes, M. Crumiller, N. M. Bouvier, D. Sachs, and A. Garcia-Sastre. Influenza a virus transmission bottlenecks are defined by infection route and recipient host. Cell, Host, and Microbe, 16:691–700, 2014.

[42] Rebecca Frise, Konrad Bradley, Neeltje Van Doremalen, Monica Galiano, Ruth A Elderfield, Peter Stilwell, Jonathan W Ashcroft, Mirian Fernandez-Alonso, Shahjahan Miah, Angie Lackenby, et al. Contact transmission of influenza virus between ferrets imposes a looser bottleneck than respiratory droplet transmission allowing propagation of antiviral resistance. Scientific Reports, 6:29793, 2016.

[43] L.H. Moncla, G. Zhong, C.W. Nelson, J.M. Dinis, J. Mutschler, A.L. Hughes, T. Watanabe, Y. Kawaoka, and T.C. Friedrich. Selective bottlenecks shape evolutionary pathways taken during mammalian adaptation of a 1918-like avian influenza virus. Cell, Host, and Microbe, 19:169–180, 2016.

[44] B. F. Keele, E. E. Giorgi, J. F. Salazar-Gonzalez, J. M. Decker, K. T. Pham, M. G. Salazar, C. Sun, T. Grayson, S. Wang, and H. Li. Identification and characterization of transmitted and early founder virus envelopes in primary HIV-1 infection. Proceedings of the National Academy of Sciences, 105:7552–7557, 2008.

[45] Damien C Tully, Colin B Ogilvie, Rebecca E Batorsky, David J Bean, Karen A Power, Musie Ghebremichael, Hunter E Bedard, Adrianne D Gladden, Aaron M Seese, Molly A Amero, et al. Differences in the selection bottleneck between modes of sexual transmission influence the genetic composition of the hiv-1 founder virus. PLoS pathogens, 12(5):e1005619, 2016.

[46] Samuel Mundia Kariuki, Philippe Selhorst, Kevin K Ariën, and Jeffrey R Dorfman. The hiv-1 transmission bottleneck. Retrovirology, 14(1):22, 2017.

[47] G. P. Wang, S. A. Sherrill-Mix, K. M. Chang, C. Quince, and F. D. Bushman. Hepatitis C virus transmission bottlenecks analyzed by deep sequencing. Journal of Virology, 84: 6218–6228, 2010.

[48] R. A. Bull, F. Luciani, K. McElroy, S. Gaudieri, S. T. Pham, A. Chopra, B. Cameron, L. Maher, G. J. Dore, and P. A. White. Sequential bottlenecks drive viral evolution in early acute hepatitis C virus infection. PLOS Pathogens, 7:e1002243., 2011.

[49] P. R. Wilker, J. M. Dinis, G. Starrett, M. Imai, M. Hatta, C. W. Nelson, D. H. O’Connor, A. L. Hughes, G. Neumann, and Y. Kawaoka. Selection on haemagglutinin imposes a bottleneck during mammalian transmission of reassortant H5n1 influenza viruses. Nature Communications, 4, 2013.

[50] John T McCrone, Robert J Woods, Emily T Martin, Ryan E Malosh, Arnold S Monto, and Adam S Lauring. Stochastic processes constrain the within and between host evolution of influenza virus. eLife, 7:e35962, 2018.

[51] Andrew L. Valesano, William J. Fitzsimmons, John T. McCrone, Joshua G. Petrie, Arnold S. Monto, Emily T. Martin, and Adam S. Lauring. Influenza b viruses exhibit lower within-host diversity than influenza a viruses in human hosts. Journal of Virology, 2019. ISSN 0022-538X. doi: 10.1128/JVI.01710-19. URL https://jvi.asm.org/content/early/2019/11/28/JVI.01710-19.

[52] P. R. Murcia, J. Hughes, P. Battista, L. Lloyd, G. J. Baillie, R. H. Ramirez-Gonzalez, D. Ormond, K. Oliver, D. Elton, and J. A. Mumford. Evolution of an Eurasian avian-like influenza virus in naive and vaccinated pigs. PLOS Pathogens, 8:e1002730, 2012.

[53] K.J. Emmett, A. Lee, H. Khiabanian, and R. Rabadan. High-resolution genomic surveillance of 2014 ebolavirus using shared subclonal variants. PLOS Currents Outbreaks, 1, 2015.

[54] L.L.M. Poon, T. Song, R. Rosenfeld, X. Lin, M.B. Rogers, B. Zhou, R. Sebra, R.A. Halpin, Y. Guan, and A. Twaddle. Quantifying influenza virus diversity and transmission in humans. Nature Genetics, 48:195–200, 2016.

[55] Ashley Sobel Leonard, Daniel Weissman, Benjamin Greenbaum, Elodie Ghedin, and Katia Koelle. Transmission bottleneck size estimation from pathogen deep-sequencing data, with an application to human influenza a virus. Journal of virology, pages JVI–00171, 2017.

[56] Katherine S Xue and Jesse D Bloom. Reconciling disparate estimates of viral genetic diversity during human influenza infections. Nature genetics, page 1, 2019.

[57] Debrah I Boeras, Peter T Hraber, Mackenzie Hurlston, Tammy Evans-Strickfaden, Tanmoy Bhattacharya, Elena E Giorgi, Joseph Mulenga, Etienne Karita, Bette T Korber, Susan Allen, et al. Role of donor genital tract hiv-1 diversity in the transmission bottleneck. Proceedings of the National Academy of Sciences, 108(46):E1156–E1163, 2011.

[58] J. M. Carlson, M. Schaefer, D. C. Monaco, R. Batorsky, D. T. Claiborne, J. Prince, M. J. Deymier, Z. S. Ende, N. R. Klatt, and C. E. DeZiel. Selection bias at the heterosexual HIV-1 transmission bottleneck. Science, 345:1254031, 2014.

[59] A. C. Hurt, S. S. Nor’e, J. M. McCaw, H. R. Fryer, J. Mosse, A. R. McLean, and I. G. Barr. Assessing the viral fitness of oseltamivir-resistant influenza viruses in ferrets using a competitive-mixtures model. Journal of Virology, 84:9427–9438, 2010.

[60] J.M. McCaw, N. Arinaminpathy, A.C. Hurt, J. McVernon, and A.R. McLean. A mathematical framework for estimating pathogen transmission fitness and inoculum size using data from a competitive mixtures animal model. PLOS Computational Biology, 7: e1002026, 2011.

[61] M. Imai, T. Watanabe, M. Hatta, S. C. Das, M. Ozawa, K. Shinya, G. Zhong, A. Hanson, H. Katsura, S. Watanabe, C. Li, E. Kawakami, S. Yamada, M. Kiso, Y. Suzuki, E. A. Maher, G. Neumann, and Y. Kawaoka. Experimental adaptation of an influenza H5 HA confers respiratory droplet transmission to a reassortant H5 HA/h1n1 virus in ferrets. Nature, 486:420–428, 2012.

[62] H. Zaraket, T. Baranovich, B.S. Kaplan, R. Carter, M. Song, J.C. Paulson, J.E. Rehg, J. Bahl, J.C. Crumpton, and J. Seiler. Mammalian adaptation of influenza a (h7n9) virus is limited by a narrow genetic bottleneck. Nature Communications, 6:6553, 2015.

[63] G. M. Shaw and E. Hunter. HIV transmission. Cold Spring Harbor perspectives in medicine, 2:a006965, 2012.

[64] S. Alizon and C. Fraser. Within-host and between-host evolutionary rates across the HIV-1 genome. Retrovirology, 10, 2013.

[65] K.A. Lythgoe, L. Pellis, and C. Fraser. Is hiv short-sighted? insights from a multistrain nested model. Evolution, 67:2769–2782, 2013.

[66] E. Sorrell, H. Wan, Y. Araya, H. Song, and D.R. Perez. Minimal molecular constraints for respiratory droplet transmission of an avian–human h9n2 influenza a virus. Proceedings of the National Academy of Sciences, 106:7565–7570, 2009.

[67] C.D. Goodman, J.E. Siregar, V. Mollard, J. Vega-Rodríguez, D. Syafruddin, H. Matsuoka, M. Matsuzaki, T. Toyama, A. Sturm, and A. Cozijnsen. Parasites resistant to the antimalarial atovaquone fail to transmit by mosquitoes. Science, 352:349–353, 2016.

[68] Marianne De Paepe and François Taddei. Viruses’life history: towards a mechanistic basis of a trade-off between survival and reproduction among phages. PLoS biology, 4(7):e193, 2006.

[69] B.R. Wasik, A. Bhushan, C.B. Ogbunugafor, and P.E. Turner. Delayed transmission selects for increased survival of vesicular stomatitis virus. Evolution, 69:117–125, 2015.

[70] Andreas Handel, Justin Brown, David Stallknecht, and Pejman Rohani. A multi-scale analysis of influenza a virus fitness trade-offs due to temperature-dependent virus persistence. PLOS Computational Biology, 9(3):1–13, 03 2013.

[71] J.R. Gog, L. Pellis, J.L.N. Wood, A.R. McLean, N. Arinaminpathy, and J.O. Lloyd-Smith. Seven challenges in modeling pathogen dynamics within-host and across scales. Epidemics, 10:45–48, 2015.

[72] M.A. Gilchrist and D. Coombs. Evolution of virulence: interdependence, constraints, and selection using nested models. Theoretical Population Biology, 69(2):145–153, 2006.

[73] Daniel Coombs, Michael A. Gilchrist, and Colleen L. Ball. Evaluating the importance of within- and between-host selection pressures on the evolution of chronic pathogens. Theoretical Population Biology, 72(4):576–591, dec 2007. doi: 10.1016/j.tpb.2007.08.005. URL https://doi.org/10.1016%2Fj.tpb.2007.08.005.

[74] N. Mideo, S. Alizon, and T. Day. Linking within-and between-host dynamics in the evolutionary epidemiology of infectious diseases. Trends in ecology and evolution, 23: 511–517, 2008.

[75] Lauren M. Childs, Fadoua El Moustaid, Zachary Gajewski, Sarah Kadelka, Ryan Nikin-Beers, Jr John W. Smith, Melody Walker, and Leah R. Johnson. Linked within-host and between-host models and data for infectious diseases: a systematic review. PeerJ, 7: e7057. jun 2019. doi: 10.7717/peerj.7057. URL https://doi.org/10.7717%2Fpeerj.7057.

[76] C. R. Parrish, E. C. Holmes, D. M. Morens, E. C. Park, D. S. Burke, C. H. Calisher, C. A. Laughlin, L. J. Saif, and P. Daszak. Cross-species virus transmission and the emergence of new epidemic diseases. Microbiology and Molecular Biology Reviews, 72:457–470, 2008.

[77] M. Woolhouse, F. Scott, Z. Hudson, R. Howey, and M. Chase-Topping. Human. viruses: discovery and emergence. Philosophical Transactions of the Royal Society B: Biological Sciences, 367:2864–2871, 2012.

[78] H. A. Orr and R. L. Unckless. The population genetics of evolutionary rescue. PLOS Genetics, 10:e1004551, 2014.

[79] T. E. Harris. The theory of branching processes. 2002. Corrected reprint of the 1963 original [Springer, Berlin].

[80] K. B. Athreya and P. E. Ney. Branching processes. Dover Publications Inc., Mineola, NY, 2004.

[81] Molly Gallagher, Christopher Brooke, Ruian Ke, and Katia Koelle. Causes and consequences of spatial within-host viral spread. Viruses, 10(11):627, 2018.

[82] Lionel B Ivashkiv and Laura T Donlin. Regulation of type i interferon responses. Nature reviews Immunology, 14(1):36, 2014.

[83] Yufan Huang, Huaiyu Dai, and Ruian Ke. Principles of effective and robust innate immune response to viral infections: a multiplex network analysis. Frontiers in immunology, 10:1736, 2019.

[84] Prasith Baccam, Catherine Beauchemin, Catherine A Macken, Frederick G Hayden, and Alan S Perelson. Kinetics of influenza a virus infection in humans. Journal of virology, 80 (15):7590–7599, 2006.

[85] Ruy M Ribeiro, Li Qin, Leslie L Chavez, Dongfeng Li, Steven G Self, and Alan S Perelson. Estimation of the initial viral growth rate and basic reproductive number during acute hiv-1 infection. Journal of virology, 84(12):6096–6102, 2010.

[86] Sebastian Bonhoeffer, Robert M May, George M Shaw, and Martin A Nowak. Virus dynamics and drug therapy. Proceedings of the National Academy of Sciences, 94(13): 6971–6976, 1997.

[87] Laetitia Canini, Jessica M Conway, Alan S Perelson, and Fabrice Carrat. Impact of different oseltamivir regimens on treating influenza a virus infection and resistance emergence: insights from a modelling study. PLoS computational biology, 10(4):e1003568, 2014.

[88] Ruian Ke, Hui Li, Shuyi Wang, Wenge Ding, Ruy M Ribeiro, Elena E Giorgi, Tanmoy Bhattacharya, Richard JO Barnard, Beatrice H Hahn, George M Shaw, et al. Superinfection and cure of infected cells as mechanisms for hepatitis c virus adaptation and persistence. Proceedings of the National Academy of Sciences, 115(30):E7139–E7148, 2018.

[89] P. Baccam, C. Beauchemin, C.A. Macken, F.G. Hayden, and A.S. Perelson. Kinetics of influenza a virus infection in humans. Journal of Virology, 80:7590–7599, 2006.

[90] Barnaby Edward Young, Sean Wei Xiang Ong, Shirin Kalimuddin, Jenny G. Low, Seow Yen Tan, Jiashen Loh, Oon-Tek Ng, Kalisvar Marimuthu, Li Wei Ang, Tze Minn Mak, Sok Kiang Lau, Danielle E. Anderson, Kian Sing Chan, Thean Yen Tan, Tong Yong Ng, Lin Cui, Zubaidah Said, Lalitha Kurupatham, Mark I-Cheng Chen, Monica Chan, Shawn Vasoo, Lin-Fa Wang, Boon Huan Tan, Raymond Tzer Pin Lin, Vernon Jian Ming Lee, Yee-Sin Leo, David Chien Lye, and for the Singapore 2019 Novel Coronavirus Outbreak Research Team. Epidemiologic Features and Clinical Course of Patients Infected With SARS-CoV-2 in Singapore. JAMA, 03 2020. ISSN 0098-7484. doi: 10.1001/jama.2020.3204. URL https://doi.org/10.1001/jama.2020.3204.

[91] Hui Li, Mark B Stoddard, Shuyi Wang, Lily M Blair, Elena E Giorgi, Erica H Parrish, Gerald H Learn, Peter Hraber, Paul A Goepfert, Michael S Saag, et al. Elucidation of hepatitis c virus transmission and early diversification by single genome sequencing. PLoS Pathogens, 8:e1002880, 2012.

[92] O. Diekmann and J.A.P. Heesterbeek. Mathematical epidemiology of infectious diseases, volume 146. Wiley, Chichester, 2000.

[93] A. Fusaro, L. Tassoni, A. Milani, J. Hughes, A. Salviato, P.R. Murcia, P. Massi, G. Zamperin, L. Bonfanti, and S. Marangon. Unexpected inter-farm transmission dynamics during a highly pathogenic avian influenza epidemic. Journal of Virology, 90:6401–11, 2016.

[94] J.M. Dinis, N.W. Florek, O.O. Fatola, L.H. Moncla, J.P. Mutschler, O.K. Charlier, J.K. Meece, E.A. Belongia, and T.C. Friedrich. Deep sequencing reveals potential antigenic variants at low frequencies in influenza a virus-infected humans. Journal of Virology, 90: 3355–3365, 2016.

[95] K. M. Pepin, S. Lass, J. R. C. Pulliam, A. F. Read, and J. O. Lloyd-Smith. Identifying genetic markers of adaptation for surveillance of viral host jumps. Nature Reviews Microbiology, 8:802–813, 2010.

[96] Katrina A Lythgoe, Andy Gardner, Oliver G Pybus, and Joe Grove. Short-sighted virus evolution and a germline hypothesis for chronic viral infections. Trends in microbiology, 25 (5):336–348, 2017.

[97] C. A. Russell, J. M. Fonville, A. E. Brown, D. F. Burke, D. L. Smith, S. L. James, S. Herfst, S. van Boheemen, M. Linster, and E. J. Schrauwen. The potential for respiratory droplet transmissible A/h5n1 influenza virus to evolve in a mammalian host. Science, 336: 1541–1547, 2012.

[98] C.T. Davis, C. Chen, L.and Pappas, J. Stevens, T.M. Tumpey, L.V Gubareva, J.M. Katz, J.M. Villanueva, R.O. Donis, and N.J. Cox. Use of highly pathogenic avian influenza A (H5N1) gain-of-function studies for molecular-based surveillance and pandemic preparedness. MBio, 5:e02431–14, 2014.

[99] T. Watanabe, G. Zhong, C.A. Russell, N. Nakajima, M. Hatta, A. Hanson, R. McBride, D.F. Burke, K. Takahashi, and S. Fukuyama. Circulating avian influenza viruses closely related to the 1918 virus have pandemic potential. Cell, Host, and Microbe, 15:692–705, 2014.

[100] Dennis Carroll, Peter Daszak, Nathan D Wolfe, George F Gao, Carlos M Morel, Subhash Morzaria, Ariel Pablos-Méndez, Oyewale Tomori, and Jonna AK Mazet. The global virome project. Science, 359(6378):872–874, 2018.

[101] C.A. Russell, P.M. Kasson, R.O. Donis, S. Riley, J. Dunbar, A. Rambaut, J. Asher, S. Burke, C.T. Davis, R.J. Garten, et al. Improving pandemic influenza risk assessment. eLife, 3:e03883, 2014.

[102] J.L. Geoghegan, A.M. Senior, F. Di Giallonardo, and E.C. Holmes. Virological factors that increase the transmissibility of emerging human viruses. Proceedings of the National Academy of Sciences, 113:4170–4175, 2016.

[103] Marc Lipsitch, Wendy Barclay, Rahul Raman, Charles J Russell, Jessica A Belser, Sarah Cobey, Peter M Kasson, James O Lloyd-Smith, Sebastian Maurer-Stroh, Steven Riley, et al. Science forum: Viral factors in influenza pandemic risk assessment. Elife, 5:e18491, 2016.

[104] M. Anishchenko, R. A. Bowen, S. Paessler, L. Austgen, I. P. Greene, and S. C. Weaver. Venezuelan encephalitis emergence mediated by a phylogenetically predicted viral mutation. Proceedings of the National Academy of Sciences of the United States of America, 103: 4994–4999, 2006.

[105] N.C. Wu, L. Dai, C.A. Olson, J.O. Lloyd-Smith, and R. Sun. Adaptation in protein fitness landscapes is facilitated by indirect paths. eLife, 5:e16965, 2016.

[106] C. Loverdo and J.O. Lloyd-Smith. Evolutionary invasion and escape in the presence of deleterious mutations. PLoS ONE, 8:e68179, 2013.

[107] Rustom Antia, Bruce R Levin, and Robert M May. Within-host population dynamics and the evolution and maintenance of microparasite virulence. The American Naturalist, 144(3): 457–472, 1994.

[108] N. Strelkowa and M. Lässig. Clonal interference in the evolution of influenza. Genetics, 192: 671–682, 2012.

[109] P.J. Gerrish and R.E. Lenski. The fate of competing beneficial mutations in an asexual population. Genetica, 102:127–144, 1998.

[110] B.H. Good, I.M. Rouzine, D.J. Balick, O. Hallatschek, and M.M. Desai. Distribution of fixed beneficial mutations and the rate of adaptation in asexual populations. Proceedings of the National Academy of Sciences, 109:4950–4955, 2012.

[111] C Brandon Ogbunugafor and Margaret J Eppstein. Competition along trajectories governs adaptation rates towards antimicrobial resistance. Nature ecology & evolution, 1(1):0007, 2017.

[112] R. Ke, J. Aaskov, E. C. Holmes, and J. O. Lloyd-Smith. Phylodynamic analysis of the emergence and epidemiological impact of transmissible defective dengue viruses. PLOS Pathogens, 9:e1003193, 2013.

